# Protocol for Membrane Permeability Prediction of Cyclic Peptides using Descriptors Obtained from Extended Ensemble Molecular Dynamics Simulations and Chemical Structures

**DOI:** 10.1101/2025.06.18.660352

**Authors:** Masatake Sugita, Yudai Noso, Jianan Li, Takuya Fujie, Keisuke Yanagisawa, Yutaka Akiyama

## Abstract

Improving membrane permeability is crucial in cyclic peptide drug discovery. Although the approach based on molecular dynamics (MD) simulation is widely used, it is computationally expensive. Alternatively, machine learning can predict membrane permeability at negligible cost, but it requires a larger dataset. There are only 7991 experimental values of membrane permeability available at the newly developed database. Another challenge in predicting membrane permeability using machine learning arises from the unique stable conformation of each cyclic peptide, which is strongly related to membrane permeability but difficult to predict from chemical structure. Therefore, we developed a machine learning protocol using 3D descriptors obtained from MD simulations in addition to 2D descriptors obtained from chemical structure of cyclic peptides, respectively, to generate a universal model with a realistic computational cost. We targeted 252 peptides across four datasets and, to calculate their 3D descriptors, predicted their conformation outside the membrane, at the water/membrane interface, and in the membrane by MD simulations based on the replica exchange with solute tempering/replica exchange umbrella sampling method using 16 replicas. For machine learning, six different algorithms were used, ranging from simple methods such as ridge regression to more sophisticated methods such as XGBoost. The best prediction performance was obtained using XGBoost, with a Pearson’s correlation coefficient *R =* 0.77 and root mean square error (*RMSE*) = 0.62. The important descriptors included those that describe the hydrophilicity and hydrophobicity of the peptide, conformational differences between inside and outside the membrane, and the degree of freedom of the peptide. We confirm the model’s ability to predict the membrane permeability of peptides that differ in chemical structure from the training data by predicting the external data consisting of 24 peptides and obtained *R* = 0.76, RMSE = 1.14. Furthermore, we extracted one of the four datasets of the training data, re-trained the model, and performed the prediction of the permeability coefficients of the extracted dataset. The results showed the model’s generic nature with *R =* 0.61 and *RMSE* = 0.74, when using position-specific (PS) 3D, and 2D descriptors. In such situations descriptors based on conformations obtained from MD are essential for the prediction.

## INTRODUCTION

Cyclic peptides, with molecular weights between those of small molecule drugs and antibody drugs, are gaining attention as a promising new pharmaceutical modality that offers the advantages of both. ^1,2^ Their larger interaction surfaces, compared to small molecule drugs, enable strong and specific binding to targets, including the inhibition of protein–protein interactions (PPIs).^3^ Their smaller molecular weight compared to antibody drugs enables oral administration and access to intracellular targets.^4^ Additionally, cyclic peptides exhibit higher metabolic stability compared to linear peptides.^5^ Consequently, cyclic peptides potentially develop drugs with higher efficacy, fewer side effects, and the ability to access previously undruggable targets for drug discovery. Furthermore, the advancement of de novo screening techniques has enabled the screening of over 10^12^ sequences, including the use of non-standard peptides.^6,7^ Techniques for rationally designing peptides based on computational methods have begun to be reported.^8,9^ With the advantages of these cyclic peptides and the development of design, synthesis, and screening techniques, a number of peptide drugs have been recently approved by the Food and Drug Administration, including two that target intracellular protein.^2^

High membrane permeability is a prerequisite for oral administration and accessing intracellular targets. However, cyclic peptides generally have low membrane permeability. Moreover, no established method for designing peptides that combine high membrane permeability with high affinity for a target has been developed. Therefore, a specific criterion for obtaining highly membrane-permeable peptides is in demand. To this end, researchers are actively investigating membrane permeation mechanisms and methods to predict membrane permeability. ^8,10–20^

Numerous experimental studies have been conducted to investigate the mechanism of membrane permeation through passive diffusion. ^10–17^ In this context, several aspects have been identified. For example, membrane permeability has a strong correlation with descriptors of hydrophilicity,^21,22^ such as AlogP.^23^ This suggests that the primary rate-limiting factor for membrane permeability is the difficulty associated with the dehydration of peptides.^24^ However, cyclic peptides can adopt unique three-dimensional (3D) structures that enable the formation of intramolecular hydrogen bonding networks, shielding their hydrophilic groups from the surroundings.^25,26^ Consequently, the membrane permeability coefficient can be significantly different even if the predicted logP value is the same, depending on the extent of shielding.^22,27^ Specifically, such chameleonic behavior is considered important for achieving both high solubility and membrane permeability, particularly for peptides with more than 10 residues.^19^ However, it is difficult to predict the 3D conformation of cyclic peptides only from their chemical structure information, making it difficult to identify membrane-permeable peptides.

In addition to experimental studies,^10–17^ computational studies have been conducted.^8,18–20^ Most computer-based analyses in this field are performed using molecular dynamics (MD) simulations.^24,28–32^ Several studies have predicted membrane permeability^24,28^ and analyzed the behavior of cyclic peptides in the membrane.^29–32^ However, accurate prediction of membrane permeability using MD simulations is computationally expensive because of the high degrees of freedom of cyclic peptides and the difficulty of sampling the cis-trans isomerization of proline and modified peptide bonds.^24^

Alternatively, membrane permeability can be predicted at a reduced cost when using machine learning methods. The newly developed database with experimental values of membrane permeability reported by Li et al., CycPeptMPDB, ^33^ has enabled the development of machine learning-based methods for predicting membrane permeability by using the dataset with 7991 instances. ^34–37^ However, the count of amino acid monomers recorded in the database is 385 types, which is sufficient to synthesize peptides comparable to 385^10^/10 types of cyclic peptides using a 10-residue peptide as an example. Furthermore, even the conversion of a few amino acids to enantiomers dramatically changes the 3D structure of cyclic peptides, resulting in a significant change in membrane permeability.^27^ Therefore, the number of experimental values reported at this time may not be sufficient to create a prediction model for membrane permeability coefficient of cyclic peptides with high generalization performance without 3D structural information inside and outside the membrane.^36^ Obviously, models developed for small molecule^38–41^ cannot be used without further consideration of uncertainty in the structure of cyclic peptides.

In this study, we developed a machine learning protocol using 3D and 2D descriptors. The 3D descriptors are obtained by predicting the peptide conformations at the outer membrane, membrane/water interface, and membrane center using MD simulations based on the replica exchange with solute tempering/replica exchange umbrella sampling method using 16 replicas, and the 2D descriptors are obtained from the chemical structure of the peptide. We targeted 252 peptides across four datasets. Six learning algorithms such as random forest and support vector machine were tested. The best prediction performance was obtained using XGBoost, with a Pearson’s correlation coefficient *R =* 0.76 and root mean square error (*RMSE*) = 0.61. The important descriptors included those that describe the hydrophilicity and hydrophobicity of the peptide, molecular weight, conformational differences between inside and outside the membrane, the degree of freedom of the peptide. We confirmed in two different ways that this model can predict membrane permeability for novel targets not included in the training data. At first, we performed predictions on external data consisting of 24 peptides and obtained *R =* 0.76, *RMSE =* 1.14. Second, we extracted one of the four datasets of the training data, re-trained the model, and predicted the permeability coefficients of the peptides of extracted dataset. The combination of descriptors that produces stably good prediction results is 2D and PS3D descriptors, and the average value of the correlation coefficient was *R =* 0.61 and *RMSE* = 0.74. In this situation, descriptors based on conformations obtained from MD are essential for the prediction.

## MATERIALS AND METHODS

### Datasets

We constructed five datasets comprising a total of 276 cyclic peptides with unique chemical structures, selected from five articles in which membrane permeability of cyclic peptides was measured using the parallel artificial membrane permeability assay (PAMPA),^42,43^ a method that utilizes artificial membranes. A summary of these datasets is presented in Table S1. The five datasets are named as Furukawa2016, Furukawa2020, Kelly2021, Bhardwaj2022, and Wang 2021, based on the first author’s name and year of publication. ^8,22,27,44,45^

Furukawa2016, Kelly2021, and Bhardwaj2022 datasets were derived from three studies that reported large numbers of data points (688, 1519, and 136, respectively) collected under uniform experimental condition. At the time this study was initiated, no other peer-reviewed literature registered in CycPeptMPDB ^33^ contained more than 100 data within a single article. The Furukawa2020 and Wang2021 datasets were selected for other reasons. We carefully selected representative peptides for each dataset. The detailed criteria for dataset selection are as follows.

#### Furukawa2016

A total of 688 membrane permeability coefficients of peptides were reported in the literature in 2016 by Furukawa et al. ^22^ In their work, two families of peptides, Library 1 and Library 2, were synthesized, and we selected Library 1 which has a common main chain structure. In this study, we only deal with cyclic peptides with an AlogP^23^ value, predicted lipophilicity, of less than 4 because membrane permeability of cyclic peptides with too high lipophilicity is considered to be not suitable for the prediction method with MD simulations.^28^ As a result, 67 of the 400 peptides in Library 1 were included as targets for this study.

#### Furukawa2020

A total of 36 membrane permeability coefficients of peptides were reported in the literature in 2020 by Furukawa et al.^44^ Among them, we selected 18 cyclic peptides (3 from library A and 15 from library B) which were used in the MD simulation by Sugita et al.^24^

#### Kelly2021

A total of 1519 membrane permeability coefficients of peptides were reported in the literature in 2021 by Kelly et al. ^27^ While a number of branched lariat-type peptides were synthesized in the literature by Kelly et al., this study targeted peptides that are non-lariat-type peptides with an AlogP value of less than 4. As a result, 86 peptides were selected as targets for this study.

#### Bhardwaj2022

A total of 136 membrane permeability coefficients of peptides were reported in the literature in 2022 by Bhardwaj et al. ^8^ In the literature, Bhardwaj et al. attempted to design peptides that could shield hydrogen bond donors from solvents by forming intramolecular hydrogen bonds in hydrophobic solvents, and they attempted to design peptides with a single stable state and peptides with multiple stable states. In this study we targeted 81 peptides with a single stable state that have an AlogP value of less than 4.

#### Wang2021

Additionally, we employed the experimental values and chemical structures of 24 peptides synthesized by Wang et al.^45^ as external test data. This dataset was not used for training.

Wang2021 was selected as test data to assess whether the model was over-learned, based on the following conditions.

1) Synthesis and assay experiments were performed by a different laboratory than the group that measured the experimental data comprising the training data.
2) The chemical structure was sufficiently similar to those in the training data.

In Bhardwaj2022 and Wang2021 datasets, the experimental values for peptides of which membrane permeability coefficients were not experimentally obtained were treated as 10^-9^ cm/s and 10^-7^ cm/s, respectively. These numbers were obtained by rounding down to the decimal point when the smallest membrane permeability value X in the respective dataset was substituted into log_10_(X). The distribution of the number of residues, molecular weight, and experimental membrane permeability (log scaled) in each dataset is shown in Figure S1. The peptides in these data sets are designed to be uncharged around pH = 7 to facilitate membrane permeation. Therefore, hydrogens and partial charge are assigned to all peptides as uncharged peptides.

## SIMULATIONS

Previous studies have suggested that the ability of cyclic peptides to adopt a closed conformation in the membrane is crucial for high membrane permeability.^18,19,31,44,46,47^ In addition, several simulation-based studies have suggested that molecular features near the water/membrane interface are important for predicting membrane permeability.^13,14,28^ Furthermore, the characteristics of peptides outside the membrane help determine whether peptides are chameleonic. Therefore, in this study, we obtained a set of peptide conformations near the membrane center, near the water/membrane interface, and outside the membrane based on MD simulations. The membrane model used was 1-palmitoyl-2-oleoyl-sn-glycero-3-phosphocholine (POPC) containing 40 mol% cholesterol, which was demonstrated by Sugita et al. to be optimal for predicting membrane permeability when using the force field parameter set described later.^24^ Specifically, 72 molecules of POPC and 48 molecules of cholesterol were used to construct the membrane model. The number of Na+ and Cl-ions was set to 16, and the number of water molecules was set to 7200 molecules. The simulation box, consisting of molecules (excluding peptide), was prepared using the Charmm-GUI server,^48^ without an additional equilibration process. This model is confirmed that peptides are fully dehydrated at the membrane center. The peptide’s location relative to the membrane was determined based on the reaction coordinate z, defined as the component orthogonal to the membrane plane of the distance between the center of mass of all nitrogen atoms of phosphatidylcholine of the membrane and that comprising the peptide bonds in the peptide main chain.

Sugita et al.^24^ have demonstrated that replica exchange with solute tempering (REST) simulations^49^ with 8 replicas are sufficient to explore the conformation of 10 residue peptides, including cis-trans isomerization of the peptide bond. In this study, we performed two sets of REST simulations at three locations: the membrane center, water/membrane interface, and outer membrane (in water) to ensure conformational redundancy. Peptides were anchored in proximity on the reaction coordinate z using umbrella potentials and exchange coordinates with each other in the replica exchange umbrella sampling (REUS) framework.^50^ That is, in this study, we ran REST/REUS simulations^50^ that exchange 2 × 8 replicas, combining 2 umbrella potentials and 8 solute tempering (ST) replicas at the three selected locations.

In this simulation protocol, the peptide was restrained using the umbrella potential, which was parabolic rather than flat-bottomed. For the REST/REUS simulation at the membrane center, two restraint positions were used: z = 0.0 Å and z = 1.0 Å. The force constants were set to 1.5 kcal/mol/Å^2^ for the umbrella potential. The peptide restraints were positioned at 13.5 Å and 15.0 Å for the REST/REUS simulation at the water/membrane interface and at 36.0 Å and 37.5 Å for the REST/REUS simulation outside the membrane. The force constants were set to 0.5 kcal/mol/Å^2^ at z = 13.5 Å, 15.0 Å, 36.0 Å, and 37.5 Å for the umbrella potential. The non-bond interaction and dihedral energies of the peptide were scaled to reflect the temperature of the peptide in each of the eight ST replicas, which corresponds to 300, 340, 390, 455, 540, 645, 785, and 980 K, when the temperature of the system was set to 300 K. Exchanges between adjacent replicas were attempted every 10 ps using the Metropolis scheme. Exchange rates were approximately 10–30%. Langevin dynamics simulation was performed at isothermal-isobaric (NPT) ensemble with semi-isotropic pressure scaling. The pressure was controlled using a Berendsen barostat and was maintained at 1 bar. The initial structure for each REST/REUS simulation was prepared using the same procedure as described by Sugita et al. and described in the supporting information.^24^ The lipid, water, and peptide molecules were parameterized using Lipid 17 force field,^51^ TIP3P model,^52^ and Amber10: Extended Hückel Theory(EHT) parameter set in molecular operating environment (MOE) from Chemical Computing Group,^53^ respectively. Amber10: EHT parameter set can be parameterized for any non-natural peptide and has been demonstrated to accurately predict the conformation of peptides in membranes targeting cyclosporine A.^54^ The REST/REUS simulations were performed for 400 ns, and snapshots were extracted at 40 ps intervals during the final 200 ns simulation. To expedite the relaxation of the model membrane structure, the first 100 ns of MD was conducted at 350 K and then reduced to 300 K over 5 ns before beginning the next simulation. For calculating the 3D descriptors at the three locations, we used 10000 snapshots obtained from 300 K replicas at two adjacent coordinates. However, for the Furukawa2020 data, snapshots of the corresponding locations were obtained from the trajectory of Sugita et al.^24^ The cost of this simulation, 3 sets of 400ns REST/REUS simulations with 16 replicas, is approximately 6/35 of that of Sugita et al, 500ns REST/REUS simulations with 224 replicas. ^24^

In this study, MD simulations are performed for a system of approximately 25,000 atoms. The calculation speed is approximately 130 ns/day for 8 processes/GPU on NVIDIA H100 GPU and AMD EPYC 9654 CPU environment with Multi Process Service (MPS). Therefore, the computational cost for a 400 ns simulation for 16 replicas is approximately 140 GPU hours. When running this simulation on the H100 GPU, the GPU’s computational resources can be effectively utilized when 8 processes/GPU or 16 processes/GPU are employed using MPS.^55^

## DESCRIPTORS

In this study, in addition to 2D descriptors calculated from the chemical structure of a peptide alone, 3D descriptors calculated from snapshots obtained from MD trajectories were used for machine learning. 3D descriptors whose values change at the three locations were treated separately from other 3D descriptors as position-specific (PS) 3D descriptors. In addition, the differences in PS3D descriptors (outer-membrane vs. water/membrane interface, outer-membrane vs. membrane center, and water/membrane interface vs. membrane center) were used as ΔPS3D descriptors.

For the 2D descriptors, we used all the descriptors available in MOE excluding those that gave the same value for all targets.^56^ The 3D descriptors used were those available in MOE, together with those obtained using CPPTRAJ module of AmberTools20.^51^ Descriptors obtained using CPPTRAJ consisted of the interaction energy between the peptide and rest (dE_vdw, dE_elec, dE), the number of hydrogen bonds between peptide and water (inter_hbond), number of intramolecular hydrogen bonds (intra_hbond, unsatisfied_hb_donor), polar surface area (polar_surface_area), conformational entropy (entropy), conformation distribution projected into PCA space of dihedral angle (dihedral_pca_cos, dihedral_pca_emd), rough shape of the peptide derived from the principal axis of inertia (p_a_i_r), angle between the peptide and membrane (p_a_i_angle_1, p_a_i_angle_2, p_a_i_angle_3), interaction between the membrane and water (LW_contacts), and change in membrane volume due to the insertion of peptide (membrane_size). The names and descriptions of all descriptors calculated using MOE are listed in Tables S2–S12. Details of the descriptors obtained using CPPTRAJ are provided in the supporting information.

The 3D and PS3D descriptors were distinguished by the correlation coefficients between descriptors computed at the membrane center, at the water/membrane interface, and in water. Thus, the correlation coefficient of the descriptor values estimated for all peptides was calculated for the following comparison: membrane center vs. water/membrane interface, membrane center vs. in water, and water/membrane interface vs. in water. If any of the three R values satisfy R < 0.9, it is considered a PS3D descriptor; otherwise, it is considered a 3D descriptor. The PS3D descriptors include each descriptor calculated at the three locations as independent descriptors, while the 3D descriptors consist of the average of the values calculated at the three locations as a descriptor value.

Note that including the ΔPS3D descriptors in the regression problem may not be necessary, as linear and nonlinear combinations of multiple descriptors are used to predict the membrane permeability coefficient. However, we aimed to assess the contribution of the ΔPS3D descriptor in the ablation study, as many physical quantities are meaningful in their differences. Therefore, we explicitly used the ΔPS3D descriptors in addition to PS3D descriptors.

## MACHINE LEARNING

Because the dataset for this study is small, deep learning algorithm was not used. Instead, lasso regression (LASSO),^57^ ridge regression (Ridge),^58^ support vector machine (SVM),^59^ random forest (RF),^60^ XGBoost,^61^ and light-GBM^62^ were used as learning algorithms. XGBoost and light-GBM are both based on boosting techniques and widely regarded as the most effective learning algorithms.^63^ RF is based on bagging and sometimes outperforms boosting methods owing to its different performance characteristics. LASSO and Ridge are basic models for linear regression, while SVM is a basic model for nonlinear regression. Three basic models were included in this study because they have fewer hyperparameters, making them less prone to overlearning. For LASSO, Ridge, and SVM, descriptors were standardized on the training data.

Since the number of descriptors used in this study was large relative to the training data, the number of descriptors used was reduced before training. Concretely, the correlation coefficients between descriptors were calculated, and one of a descriptor pair whose correlation coefficients were higher than the threshold value was deleted. The threshold was determined by evaluating MAE obtained by performing a 5-fold cross validation using random forest algorithm on the dataset. In this procedure default hyperparameters are used. Figure S2 plots the mean and standard deviation of MAE when the threshold was changed from 0.4 to 0.95 in intervals of 0.05. Since Figure shows that there is almost no difference in the results when the threshold is increased above 0.6, we decided to use a threshold value of 0.6. The descriptors chosen were 20, 1, 28, and 45 for 2D, 3D, PS3D, and ΔPS3D, respectively, for a total of 94 descriptors.

Hyperparameters were determined based on Steps 1–4, as described below. The range of searched and selected hyperparameters are presented in Tables S13–S18.

1. The dataset was randomly divided into five nonoverlapping blocks of equal size. One block was used as the validation set, and the remaining four were used as training data.
2. A 4-fold inner cross-validation was performed for each of the five validation set selection, resulting in a total of 20 results. The average of the 20 mean absolute errors (MAEs), <MAE>_20_, between the predicted and experimental values was calculated as the performance measure.
3. Step 2 was performed for all possible hyperparameter combinations.
4. The hyperparameter set with the smallest <MAE>_20_ value was selected as the optimal hyperparameter set.

The prediction performance of the membrane permeability coefficient was evaluated as the average values obtained in the process of 5-fold cross-validation using a selected hyperparameter set. Pearson’s correlation coefficient, R, coefficient of determination (Q^2^ for validation, R^2^ for test), MAE, and RMSE are used as performance parameters. All evaluations were based on the logarithm of the membrane permeability coefficient.

Notably, the membrane permeability predicted from molecular simulations is intrinsic permeability; however, in this study, the apparent permeability obtained from experiments is learned. Therefore, this protocol predicts the apparent permeability.

## RESULTS

First, we show the prediction performance on the whole dataset in Table 1. We used six algorithms for training and prediction: LASSO, Ridge, SVM, RF, XGBoost, and Light-GBM. No significant difference exists in prediction accuracy among the algorithms, with 0.77 ≥ *R* ≥ 0.73, 0.46 ≤ *MAE* ≤ 0.51, 0.61 ≤ *RMSE* ≤ 0.67, and 0.58 ≥ *Q*^2^ ≥ 0.53. The Friedman test and post-hoc Wilcoxon signed-ranks test showed *p-values* > 0.42, which did not confirm significant differences; however, the most sophisticated algorithm employed, XGBoost, obtained the best performance with *R* = 0.77, *MAE* = 0.46, *RMSE* = 0.62, and *Q*^2^ = 0.58. Hereafter, we analyzed the prediction results using XGBoost, which provided the best prediction accuracy.

**Table 1.** Prediction performance of each algorithm.

| Algorithm | R | MAE | RMSE | Q<sup>2</sup> |
| --- | --- | --- | --- | --- |
| LASSO | $0.73 \pm 0.07$ | $0.51 \pm 0.04$ | $0.65 \pm 0.05$ | $0.53 \pm 0.10$ |
| Ridge | $0.73 \pm 0.07$ | $0.51 \pm 0.05$ | $0.65 \pm 0.05$ | $0.53 \pm 0.10$ |
| SVM | $0.74 \pm 0.06$ | $0.51 \pm 0.05$ | $0.65 \pm 0.06$ | $0.53 \pm 0.09$ |
| RF | $0.76 \pm 0.06$ | $0.48 \pm 0.06$ | $0.64 \pm 0.08$ | $0.55 \pm 0.08$ |
| XGBoost | $0.77 \pm 0.07$ | $0.46 \pm 0.05$ | $0.62 \pm 0.08$ | $0.58 \pm 0.08$ |
| LightGBM | $0.74 \pm 0.07$ | $0.47 \pm 0.06$ | $0.65 \pm 0.09$ | $0.54 \pm 0.01$ |

We compared the prediction performance of the validation for the four individual datasets. Table 2 presents the correlation coefficient R, MAE, RMSE and Q^2^ for each dataset separately. The predicted and experimental values for all validation data are plotted as scatter plots in Figure 1. Furukawa2016 has *R =* 0.86, *MAE* = 0.41, *RMSE* = 0.51, *Q*^2^ = 0.71, Kelly2021 has *R =* 0.83, *MAE* = 0.54, *RMSE* = 0.40, *Q*^2^ = 0.69; the permeability of these two datasets can be predicted with high accuracy. Bhardwaj2022 has middle accuracy, *R* = 0.70, *MAE* = 0.65, *RMSE* = 0.85, and *Q*^2^ = 0.47. Furukawa2020 is less accurate with *R* = 0.45, *MAE* = 0.54, *RMSE* = 0.66, and *Q*^2^ = 0.17. This result demonstrate that the prediction accuracy varied considerably among the datasets. Values with other algorithms are shown in Tables S19–S23.

**Table 2.** Prediction performance of XGBoost against each dataset.

| Data | R | MAE | RMSE | Q<sup>2</sup> |
| --- | --- | --- | --- | --- |
| Furukawa2016 | 0.86 | 0.41 | 0.51 | 0.71 |
| Furukawa2020 | 0.45 | 0.54 | 0.66 | 0.17 |
| Kelly2021 | 0.83 | 0.31 | 0.40 | 0.69 |
| Bhardwaj2022 | 0.70 | 0.65 | 0.85 | 0.47 |

**Figure 1.**
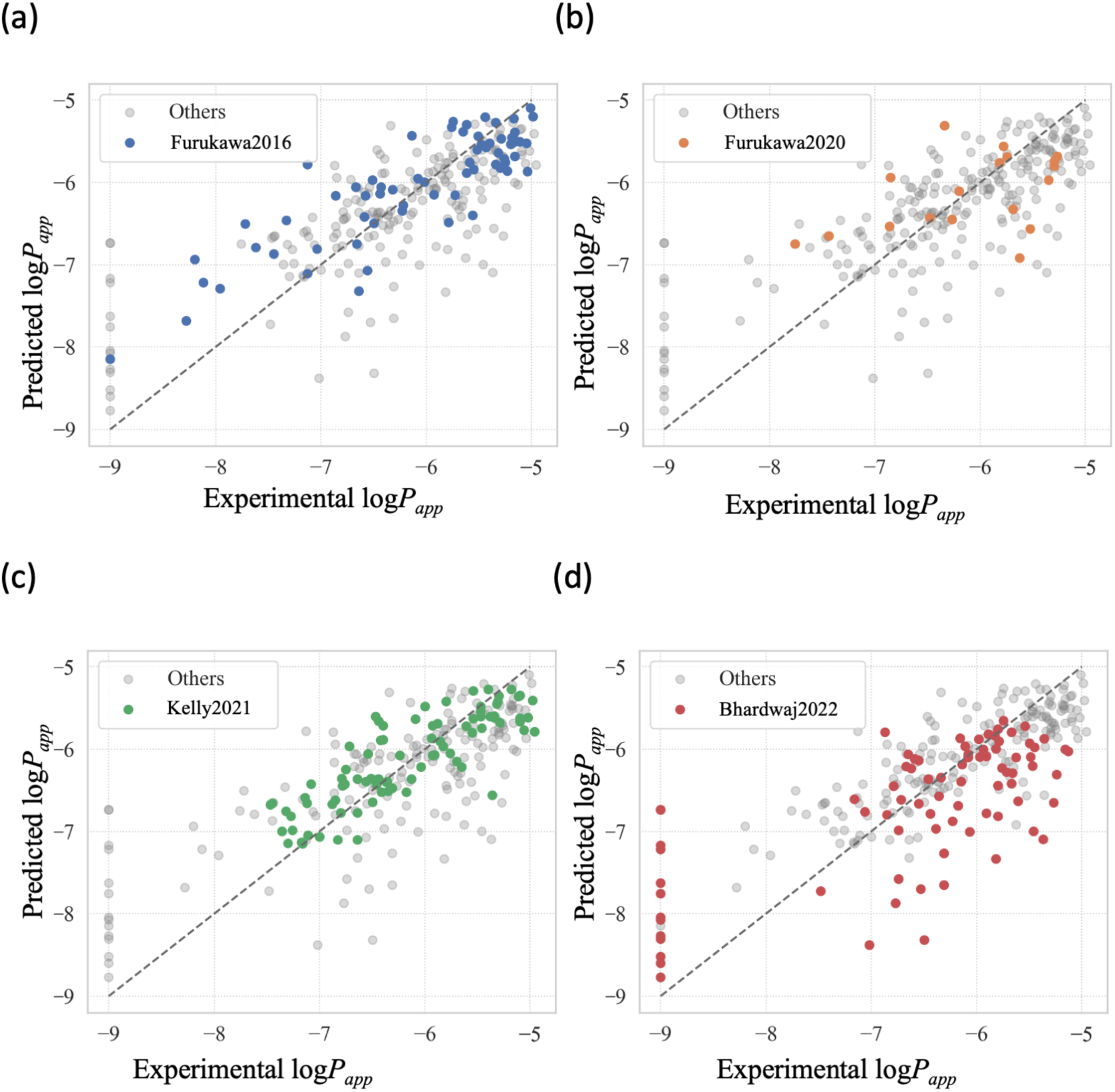
Scatter plot of predicted and experimental values for all validation data. XGBoost is chosen as the learning algorithm.

In the prediction results for Bhardwaj2022, there is a large variation in the prediction results of permeability coefficients for peptides under the experimental detection limit as shown in Figure 1. There are two possible reasons for this large variance. The first possibility is that there are peptides with membrane permeability coefficients significantly smaller than 10^-9^. In other words, the correlation with the calculated value may increase when the experimental detection limit is raised to approximately 10^-12^. The second possibility is that the source of the low membrane permeability of peptides below the detection limit may not be reproducible with the current MD setup. That is, membrane permeation of peptides that cannot be detected experimentally even though the predicted membrane permeation coefficient from MD is greater than 10^-8^ may be prevented by the effect of an unstirred water layer^43,64^ or other unknown mechanisms. The main reason for the low prediction accuracy of the Furukawa2020 dataset is likely the small sample size (18 peptides). In each five-fold cross-validation trials, approximately four peptides are used as the validation set. This limited size increases the probability that peptides with similar membrane permeability coefficients are grouped together in the validation set, which leads to a significant underestimation of the R and Q^2^ values. Furthermore, 3 of the 18 peptides were selected from Library A, whereas the remaining 15 were from Library B. Peptides from Library A differ significantly in their AlogP– permeability relationships and conformational behavior in solution.^44^ Therefore, it is expected that the prediction accuracy tends to decrease where peptides from Library A are included in the validation set.

To determine which set of descriptors (2D, 3D, PS3D, and ΔPS3D) made the most significant contribution, we again built and validated models using each set of descriptors alone and their subsets but same hyperparameters. The results are presented in Table 3. The most important contribution is made by the 2D descriptors, with *R =* 0.77, *MAE* = 0.46, *RMSE* = 0.59, and *Q*^2^ = 0.56 on the experimental data. The predictions obtained using only the PS3D descriptors have *R =* 0.57, *MAE* = 0.61, *RMSE* = 0.80, and *Q*^2^ = 0.28 showing a middle contribution to membrane permeability prediction. The prediction performance made using only ΔPS3D is *R =* 0.23, *MAE* = 0.76, *RMSE* = 0.97, and *Q*^2^ = −0.04 and those obtained using only 3D have *R =* 0.25, *MAE* = 0.83, *RMSE* = 1.15, and *Q*^2^ = −0.55, indicating a significantly low contribution to membrane permeability prediction. There were three patterns of descriptor combinations that yielded more accurate prediction results than using all descriptors (2D and PS3D descriptors, 2D, 3D, and PS3D descriptors, and 2D, PS3D, and ΔPS3D descriptors). These results demonstrate that the 2D and PS3D descriptors are important to predict the membrane permeation coefficients.

**Table 3.** Prediction performance when using all descriptors and their subsets. A + mark denotes that the corresponding descriptor was used for training and validation.

| 2D | 3D | PS3D | ΔPS3D | R | MAE | RMSE | Q<sup>2</sup> |
| --- | --- | --- | --- | --- | --- | --- | --- |
| + | + | + | + | $0.77 \pm 0.06$ | $0.46 \pm 0.05$ | $0.62 \pm 0.06$ | $0.58 \pm 0.08$ |
| | + | + | + | $0.57 \pm 0.01$ | $0.61 \pm 0.08$ | $0.80 \pm 0.11$ | $0.29 \pm 0.14$ |
| + | | + | + | $0.78 \pm 0.06$ | $0.46 \pm 0.05$ | $0.61 \pm 0.07$ | $0.58 \pm 0.08$ |
| + | + | | + | $0.75 \pm 0.08$ | $0.48 \pm 0.05$ | $0.64 \pm 0.07$ | $0.54 \pm 0.10$ |
| + | + | + | | $0.79 \pm 0.05$ | $0.44 \pm 0.05$ | $0.59 \pm 0.08$ | $0.61 \pm 0.07$ |
| | | + | + | $0.51 \pm 0.12$ | $0.63 \pm 0.08$ | $0.84 \pm 0.11$ | $0.23 \pm 0.16$ |
| | + | | + | $0.35 \pm 0.11$ | $0.72 \pm 0.07$ | $0.92 \pm 0.08$ | $0.07 \pm 0.14$ |
| | + | + | | $0.60 \pm 0.11$ | $0.60 \pm 0.08$ | $0.78 \pm 0.12$ | $0.33 \pm 0.16$ |
| + | | | + | $0.75 \pm 0.08$ | $0.48 \pm 0.05$ | $0.64 \pm 0.08$ | $0.55 \pm 0.10$ |
| + | | + | | $0.79 \pm 0.05$ | $0.44 \pm 0.05$ | $0.60 \pm 0.08$ | $0.61 \pm 0.08$ |
| + | + | | | $0.75 \pm 0.03$ | $0.49 \pm 0.03$ | $0.65 \pm 0.06$ | $0.53 \pm 0.03$ |
| | | | + | $0.23 \pm 0.14$ | $0.76 \pm 0.06$ | $0.97 \pm 0.07$ | $-0.04 \pm 0.16$ |
| | | + | | $0.57 \pm 0.15$ | $0.61 \pm 0.08$ | $0.80 \pm 0.14$ | $0.28 \pm 0.21$ |
| | + | | | $0.25 \pm 0.11$ | $0.83 \pm 0.08$ | $1.15 \pm 0.10$ | $-0.55 \pm 0.50$ |
| + | | | | $0.77 \pm 0.04$ | $0.46 \pm 0.04$ | $0.63 \pm 0.06$ | $0.56 \pm 0.04$ |

Subsequently, the importance of each descriptor was calculated in the prediction model using XGBoost. The 10 descriptors (six 2D, two ΔPS3D, one PS3D, and one 3D) with the highest contribution are shown in Figure 2. Details of these descriptors are provided in Table S24. Six descriptors—SlogP, density, GCUT_SLOGP_1, GCUT_PEOE_0, vsurf_CW5_sur, vsurf_IW1_mem - vsurf_IW1_sur—reflect hydrophilic and hydrophobic characteristics. Opr_nrot, representing the number of rotatable bonds, relates to the molecule size and conformation entropy change of the peptide. Vsa_pol denotes polar surface area but is highly correlated with the molecular weight of the peptide (R = 0.89), rather than SlogP. E_str, stretching potential, also correlates with molecular weight (R = 0.48). Dihedral_pca_cos (membrane - water) denotes the extent of the conformation change between the membrane center and water phase. Of these, SlogP and GCUT_SLOGP_1 were positively correlated with membrane permeability (R = 0.47, R = 0.44), while density and vsurf_CW5_sur were negatively correlated (R = 0.45, R = 0.41). Thus, the hydrophilic and hydrophobic features directly influence membrane permeability across the entire dataset. Additionally, other factors (e.g. molecular weight and peptide degrees of freedom) appear to determine the deviation from the trend formed by the hydrophilicity/hydrophobicity of the peptides and the membrane permeability coefficient. The high contribution of Dihedral_pca_cos (membrane - water) suggests that the extent of peptide conformational change as it enters the membrane from aqueous solution is related to its membrane permeability.

**Figure 2.**
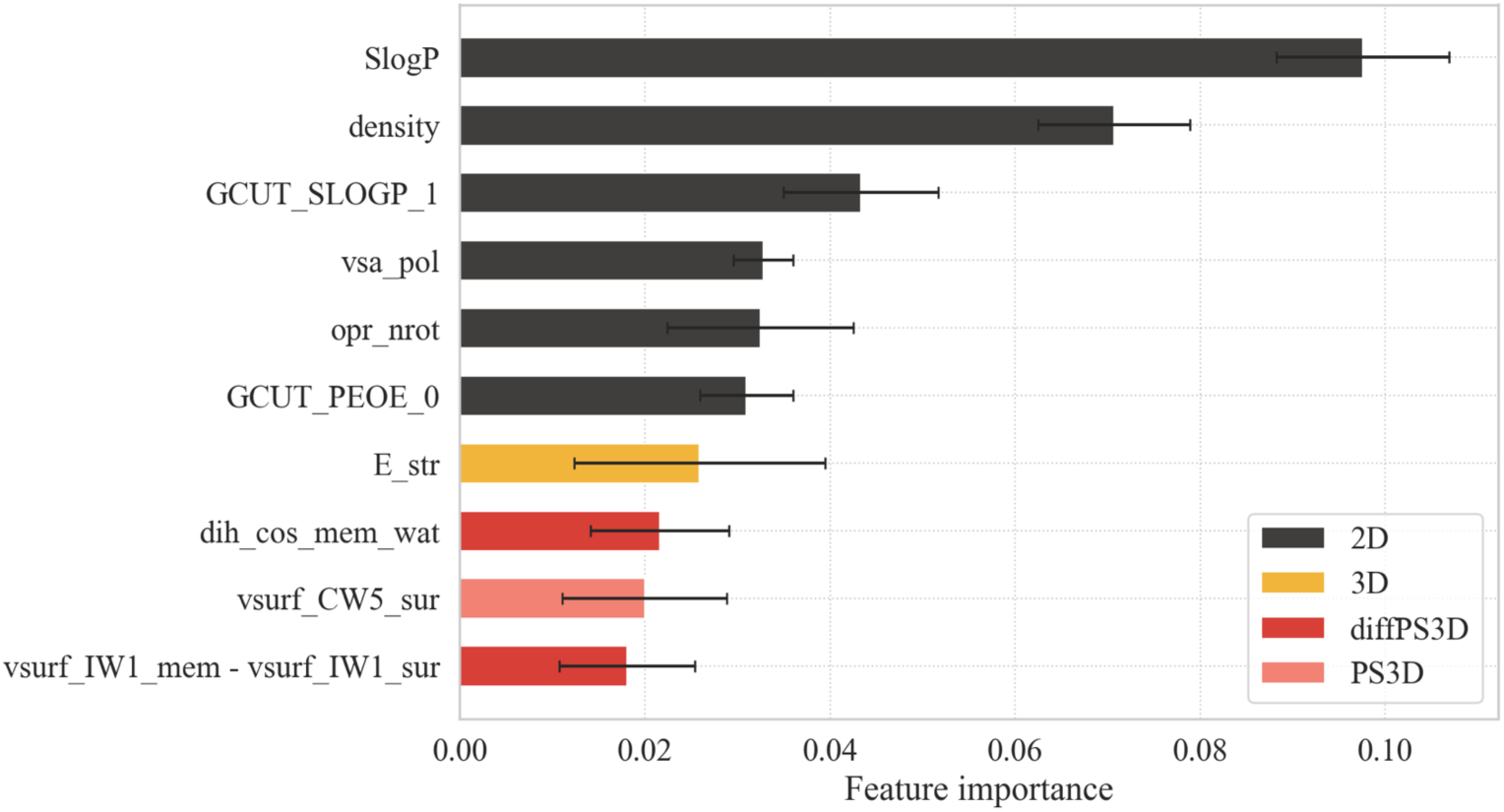
Ten most important descriptors with the highest importance.

Subsequently, we predicted membrane permeability of external data, Wang2021, which was completely independent of the training data. We performed a principal component analysis on the 94 descriptors used in the train, validation, and external test, and plotted the peptides comprising four datasets in the train and validation data, as well as the peptides in Wang2021 against the first and second principal components in Figure 3. The peptides in Wang2021 are within the descriptor space of the training data, suggesting that the model can successfully predict membrane permeability if it is not overfitted.

**Figure 3.**
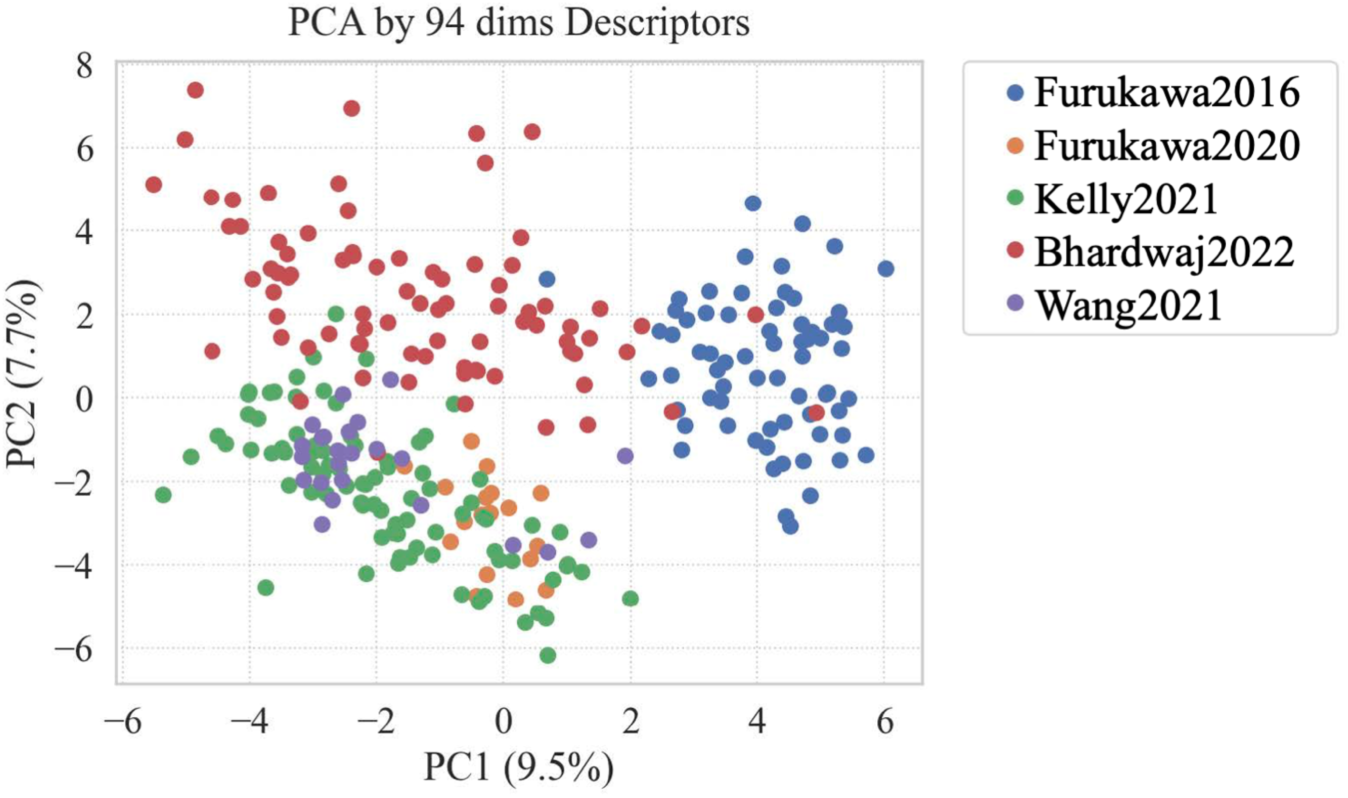
Scatter plot on the first and second axis estimated from a principal component analysis of the 2D descriptors. The number in parentheses indicate the contribution.

Experimental and predicted values of membrane permeability for the Wang2021 are plotted in Figure 4. The prediction accuracy was *R* = 0.76, *MAE* = 0.96, *RMSE* = 1.14, and *Q*^2^ = −0.06. The correlation coefficient was 0.76, which is approximately the average of the correlation coefficient R obtained in the validation process. Since the most important factor in screening is the rank of membrane permeability, the protocol proposed in this study can be used to predict new data in this respect. However, the predicted values of membrane permeability coefficients are generally lower than the experimental values, and thus the evaluation values that depend on the difference between the predicted and experimental values are low. This is because of the values of membrane permeability reported in the literature of Wang2021, which are approximately one order of magnitude higher than peptides with similar molecular weights and SlogP values in other datasets. As shown in other literature,^44^ this is caused by the differences in experimental protocols, which can lead to occur with a certain probability when predicting new data for membrane permeability. Anyway, it is very significant that we were able to obtain predicted values of membrane permeability for external data that correlate well with the experimental values.

**Figure 4.**
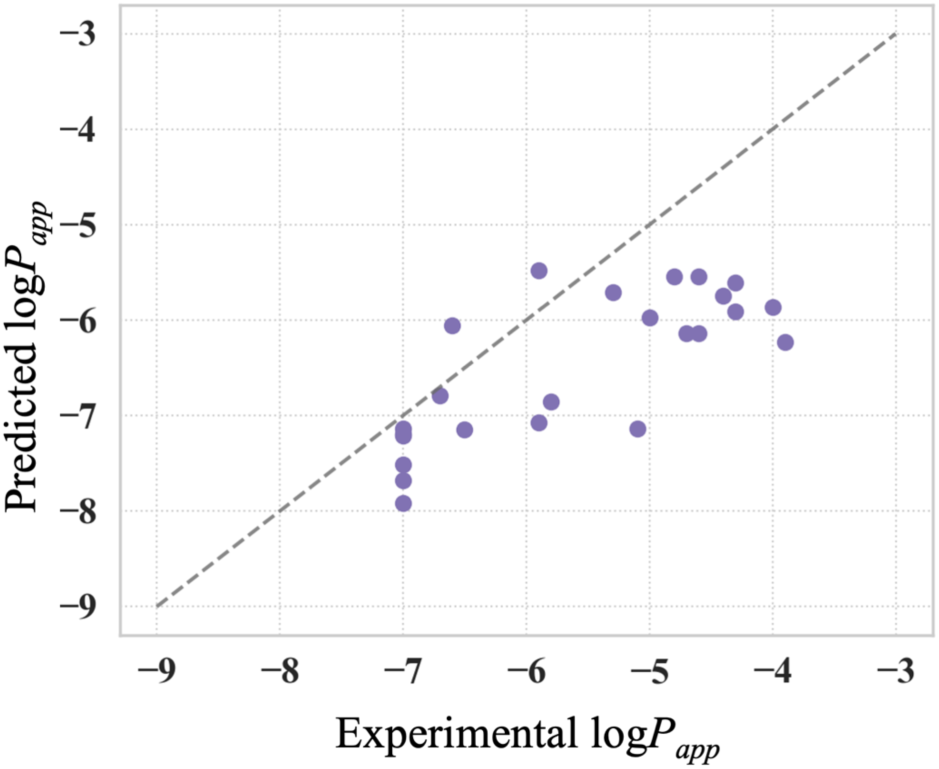
Scatter plot of predicted and experimental values for Wang2021 data. XGBoost is chosen as the learning algorithm.

The generalization performance of the method and characteristics of each dataset were further checked. That is, we used all peptides from any one of the four datasets as external test data and the remaining as training data but using the same hyperparameters presented in Table S17. Tables 4–7 show the prediction performance of Furukawa2016, Furukawa2020, Kelly2021, and Bhardwaj2022 when each was used as test data. In this calculation, the only case in which the 2D descriptors successfully predicted membrane permeability is when Furukawa2016 is used as test data (*R =* 0.79, *MAE* = 0.52, *RMSE* = 0.63, *R*^2^ = 0.57). Predicting membrane permeability using only 2D descriptors is difficult for the other datasets. Instead, PS3D descriptors have to be used to successfully predict membrane permeability; In Furukawa2020 the highest prediction performance is obtained when PS3D and 2D descriptors are used, with *R =* 0.60, *MAE* = 0.47 and *RMSE* = 0.58, and *R*^2^ *=* 0.35. In Kelly2021, the highest prediction performance is obtained when only PS3D descriptors are used, with *R =* 0.59, *MAE* = 0.49, *RMSE* = 0.58, and *R*^2^ = 0.33. In Bhardwaj2022, the highest prediction performance is obtained when 2D, 3D, and PS3D descriptors are used, with *R =* 0.60, *MAE* = 0.79, *RMSE* = 0.97, and *R*^2^ = 0.32. The most stable prediction performance is obtained when using PS3D, and 2D descriptors, with an average performance of *R =* 0.61, *MAE* = 0.61, *RMSE* = 0.74, and *R*^2^ = 0.28. These results demonstrate that the importance of PS3D descriptors for predicting the membrane permeation coefficient of external data.

**Table 4.** Prediction performance when using Furukawa2016 as test data. A + mark denotes that the corresponding descriptor was used for training and validation.

| 2D | 3D | PS3D | ΔPS3D | # Feature | R | MAE | RMSE | Q<sup>2</sup> |
| --- | --- | --- | --- | --- | --- | --- | --- | --- |
| + | + | + | + | 94 | 0.72 | 0.54 | 0.68 | 0.49 |
|  | + | + | + | 74 | 0.29 | 0.69 | 0.93 | 0.05 |
| + |  | + | + | 93 | 0.77 | 0.52 | 0.64 | 0.56 |
| + | + |  | + | 66 | 0.75 | 0.56 | 0.69 | 0.48 |
| + | + | + |  | 49 | 0.74 | 0.52 | 0.65 | 0.53 |
|  |  | + | + | 73 | 0.20 | 0.74 | 0.98 | -0.05 |
|  | + |  | + | 46 | 0.09 | 0.81 | 1.00 | -0.10 |
|  | + | + |  | 29 | 0.32 | 0.68 | 0.92 | 0.07 |
| + |  |  | + | 65 | 0.80 | 0.55 | 0.67 | 0.51 |
| + |  | + |  | 48 | 0.76 | 0.52 | 0.64 | 0.55 |
| + | + |  |  | 21 | 0.77 | 0.52 | 0.64 | 0.56 |
|  |  |  | + | 45 | 0.07 | 0.88 | 1.05 | -0.20 |
|  |  | + |  | 28 | 0.19 | 0.75 | 0.96 | -0.01 |
|  | + |  |  | 1 | 0.13 | 0.86 | 1.25 | -0.72 |
| + |  |  |  | 20 | 0.79 | 0.52 | 0.63 | 0.57 |

## DISCUSSIONS

In this study, we combined MD simulations and machine learning to predict membrane permeability. Although there were no large differences among the machine learning algorithms, XGBoost gave the best prediction performance, with *R* = 0.77, *MAE* = 0.46, *RMSE* = 0.62, and *Q*^2^ = 0.58. The prediction accuracy varied considerably among the datasets as shown in Table 2. Furukawa2016 and Kelly2021 were able to predict membrane permeability with high accuracy. Bhardwaj2022 had a middle accuracy. Furukawa2020 had a slightly lower accuracy.

One factor contributing to the difference in prediction accuracy among the datasets is the relationship between the most important descriptor, SlogP, and membrane permeability. As shown in Figure 5, the membrane permeability and SlogP for Furukawa2016 and Kelly2021 are highly correlated with *R =* 0.66 and *R =* 0.83, respectively. For Furukawa2020 and Bhardwaj2022, the correlation between SlogP and membrane permeability is *R =* 0.51 and *R =* 0.55, respectively, indicating that membrane permeability cannot be predicted with high accuracy from this descriptor alone. Another reason for the difference in prediction accuracy among the datasets is the descriptor space covered by each dataset. Figure 3 shows that Furukawa2020 and Kelly2021 are consolidated in a relatively small space in the descriptor space (PC1 and PC2), while Bhardwaj2022 is distributed over a wide area. Therefore, if a peptide on the edge of the descriptor space in the Bhardwaj2022 data is included in the validation set, its predictions are likely to be inaccurate. Figure 1(d) and the MAE and RMSE values in the Table 2 show that the predicted membrane permeability coefficients of peptides in the Bhardwaj2022 data tend to deviate from the experimental values. In the case of small data sets, such as Furukawa2020, the contribution to learning and the number of peptides in the validation set for each of five cross-validation runs are significantly small; thus, the prediction accuracy during validation becomes less accurate.

**Figure 5.**
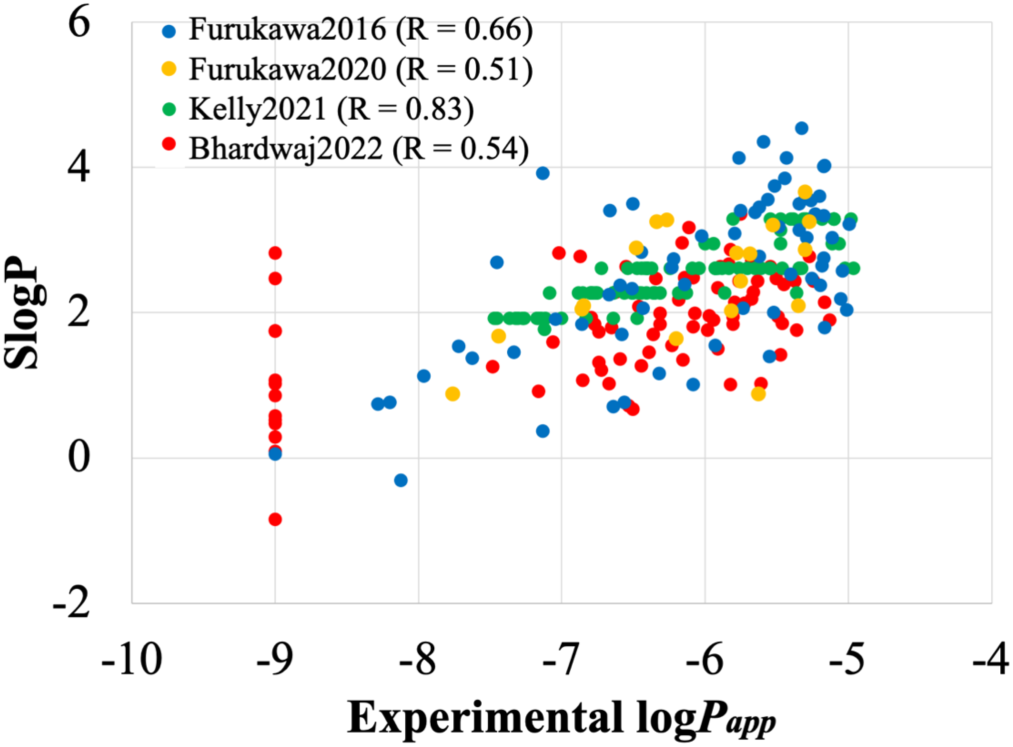
Scatter plot between SlogP and log scaled membrane permeability coefficient.

Table 3 shows the descriptors (2D, 3D, PS3D, and ΔPS3D) that made the most significant contribution to the prediction accuracy during validation. Comparing the prediction quality of each descriptor set, the 2D descriptors provided the most accurate prediction of membrane permeability during validation, where the prediction targets were randomly selected from the entire data set. The PS3D descriptors performed the second best. The high prediction performance of the 2D descriptors during validation is straightforward, as similar peptides to those in the training data are always included in the validation set. As shown in Figure 5, SlogP, the descriptor with the highest importance in validation, has a very high correlation with the logarithm of the membrane permeability coefficient; thus, a certain level of prediction may be possible by using SlogP alone. However, the combination of the 2D and PS3D descriptors has superior performance in predicting the membrane permeability coefficient. This indicates that information on peptide conformation also contributes to the prediction accuracy. The reason why the prediction accuracy is not the highest when all descriptors are used may be because the number of descriptors is numerous against the number of data (276).

When using one of the four datasets as external test data, unlike the data presented in Table 2 where the targets for the validation set were randomly selected from four datasets, the prediction accuracy using only 2D descriptors is mostly low. Only Furukawa2016 provides accurate prediction results using only 2D descriptors, but PS3D descriptors are required to predict the membrane permeability for other datasets. The average value when using 2D, and PS3D descriptors is *R =* 0.61, *MAE* = 0.61, *RMSE* = 0.74, and *R*^2^ = 0.28, which is not high, but provides some universal predictability. The results when Kelly2021 was used as test data were intriguing. The prediction performance using only 2D descriptors was *R =* 0.29, while the prediction performance using PS3D descriptors was *R =* 0.59. However, the most important 2D descriptor, SlogP, correlates well with the experimental value with a correlation coefficient of *R =* 0.83. Furukawa2016, Furukawa2020, and Bhardwaj2022 also have correlation coefficients between SlogP and experimental values of *R =* 0.66, *R =* 0.51, and *R =* 0.54, respectively. This discrepancy demonstrates the difficulty of predicting the membrane permeability of peptides having different chemical structures from the training data based only on 2D descriptors. This is because that the correlation trend between 2D descriptors such as SlogP with the permeability coefficient differs across datasets, although all datasets correlate with 2D descriptors, probably because the characteristics of their 3D structure differ. Although 2D descriptors such as SlogP describe well the trend of membrane permeability, it is expected that the corrections made by descriptors based on 3D conformation will enable more universal predictions.

As shown in Figure 2, only dihedral_pca_cos (membrane - water) was among the top 10 most important descriptors obtained from MD, indicating that descriptors obtained from MD do not appear significantly important. The primary reason why descriptors obtained from MD appear unimportant is that the membrane permeability correlate well with the 2D descriptors such as SlogP in each of dataset. For example, as shown in Table 4, when Furukawa2016 was excluded from the training data and used as test data, a correlation coefficient of *R =* 0.79 was obtained using only 2D descriptors. The correlation coefficients of *R =* 0.48 and *R* = 0.50 are also obtained for Furukawa2020 and bhardwaj2022 using only 2D descriptors as shown in Tables 5 and 7. Thus, the learned model have a higher importance of 2D descriptors, and the use of only 2D descriptors may be sufficient when predicting the membrane permeability of peptides with similar properties to the training data, such as in the validation process. The 3D descriptors may play an important role when predicting the membrane permeability of peptides with different properties from the training data, such as when predicting new data. The second reason is that various uncertainties in the physical quantities obtained from the simulations. For instance, the values obtained from the simulations strongly depend on the force field parameters and are affected by large fluctuations of potential energy. Furthermore, equations for estimating the physical quantity, such as entropy, involve a large approximation. Specifically, systems with many charged molecules, such as lipid membranes, have significantly large energetical fluctuations, and their predictions tend to have numerous statistical errors. This may be the main reason of that the ΔPS3D descriptors, which represent the difference of 3D descriptor values along the position across the membrane, reduced prediction accuracy even though the many physical quantities find meaning in the difference of states. There has been no large-scale examination into which physical quantities calculated from MD can successfully predict the membrane permeability of cyclic peptides. Furthermore, there is still room for improvement in the computational method of the PS3D and ΔPS3D descriptors used at this time, and replacing these with more effective descriptors is an important task to be addressed.

**Table 5.**
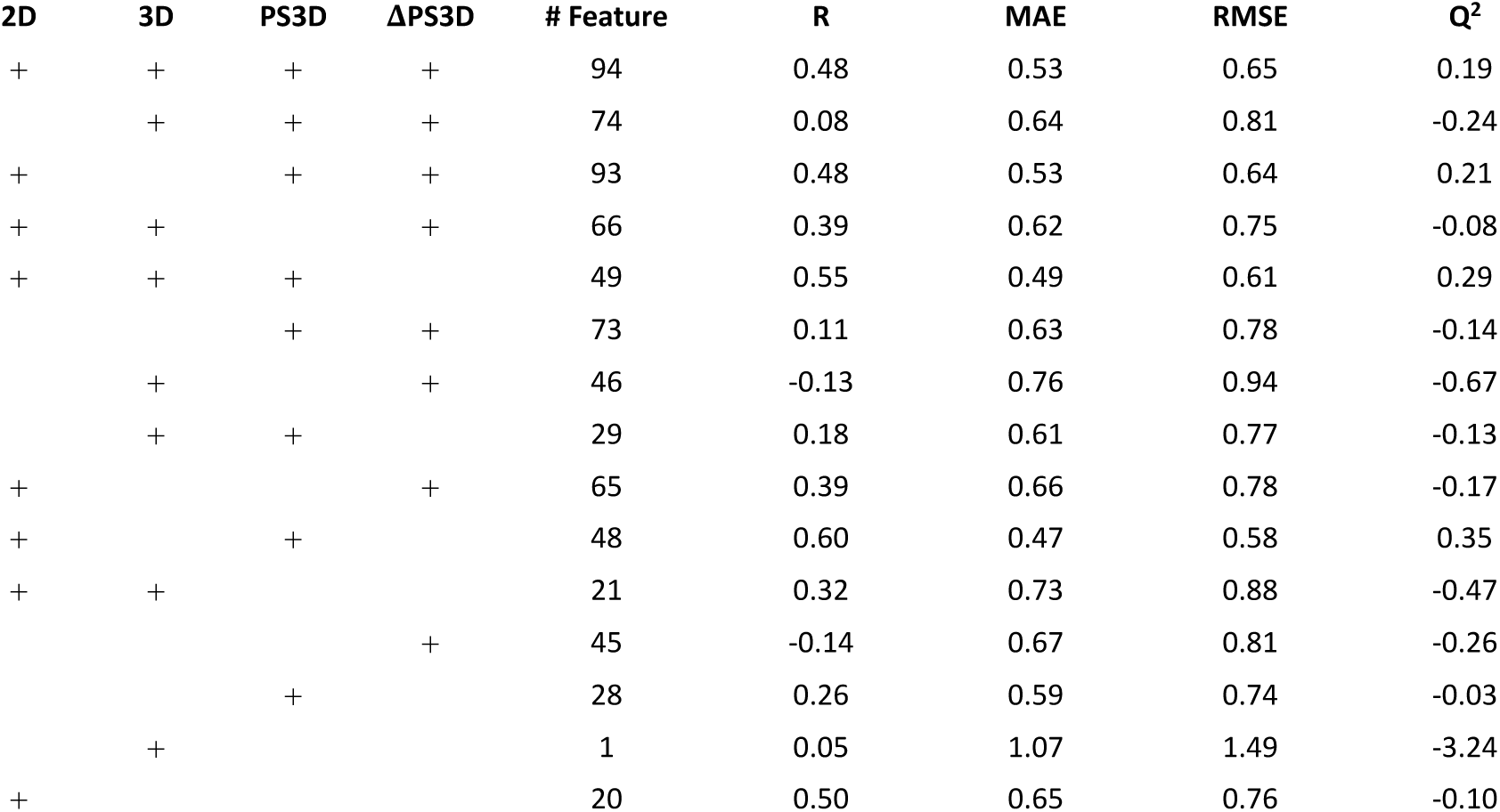
Prediction performance when using Furukawa2020 as test data. A + mark denotes that the corresponding descriptor was used for training and validation.

**Table 6.** Prediction performance when using Kelly2021 as test data. A + mark denotes that the corresponding descriptor was used for training and validation.

| 2D | 3D | PS3D | ΔPS3D | # Feature | R | MAE | RMSE | Q<sup>2</sup> |
| --- | --- | --- | --- | --- | --- | --- | --- | --- |
| + | + | + | + | 94 | 0.37 | 0.64 | 0.76 | -0.14 |
|  | + | + | + | 74 | 0.42 | 0.55 | 0.66 | 0.15 |
| + |  | + | + | 93 | 0.41 | 0.63 | 0.74 | -0.09 |
| + | + |  | + | 66 | -0.05 | 0.72 | 0.86 | -0.46 |
| + | + | + |  | 49 | 0.47 | 0.65 | 0.76 | -0.15 |
|  |  | + | + | 73 | 0.42 | 0.56 | 0.68 | 0.10 |
|  | + |  | + | 46 | -0.05 | 0.73 | 0.86 | -0.47 |
|  | + | + |  | 29 | 0.50 | 0.54 | 0.62 | 0.23 |
| + |  |  | + | 65 | -0.02 | 0.71 | 0.85 | -0.43 |
| + |  | + |  | 48 | 0.47 | 0.64 | 0.75 | -0.11 |
| + | + |  |  | 21 | 0.31 | 0.68 | 0.80 | -0.25 |
|  |  |  | + | 45 | -0.03 | 0.74 | 0.92 | -0.67 |
|  |  | + |  | 28 | 0.59 | 0.49 | 0.58 | 0.33 |
|  | + |  |  | 1 | -0.18 | 0.80 | 0.97 | -0.87 |
| + |  |  |  | 20 | 0.29 | 0.67 | 0.79 | -0.22 |

**Table 7.**
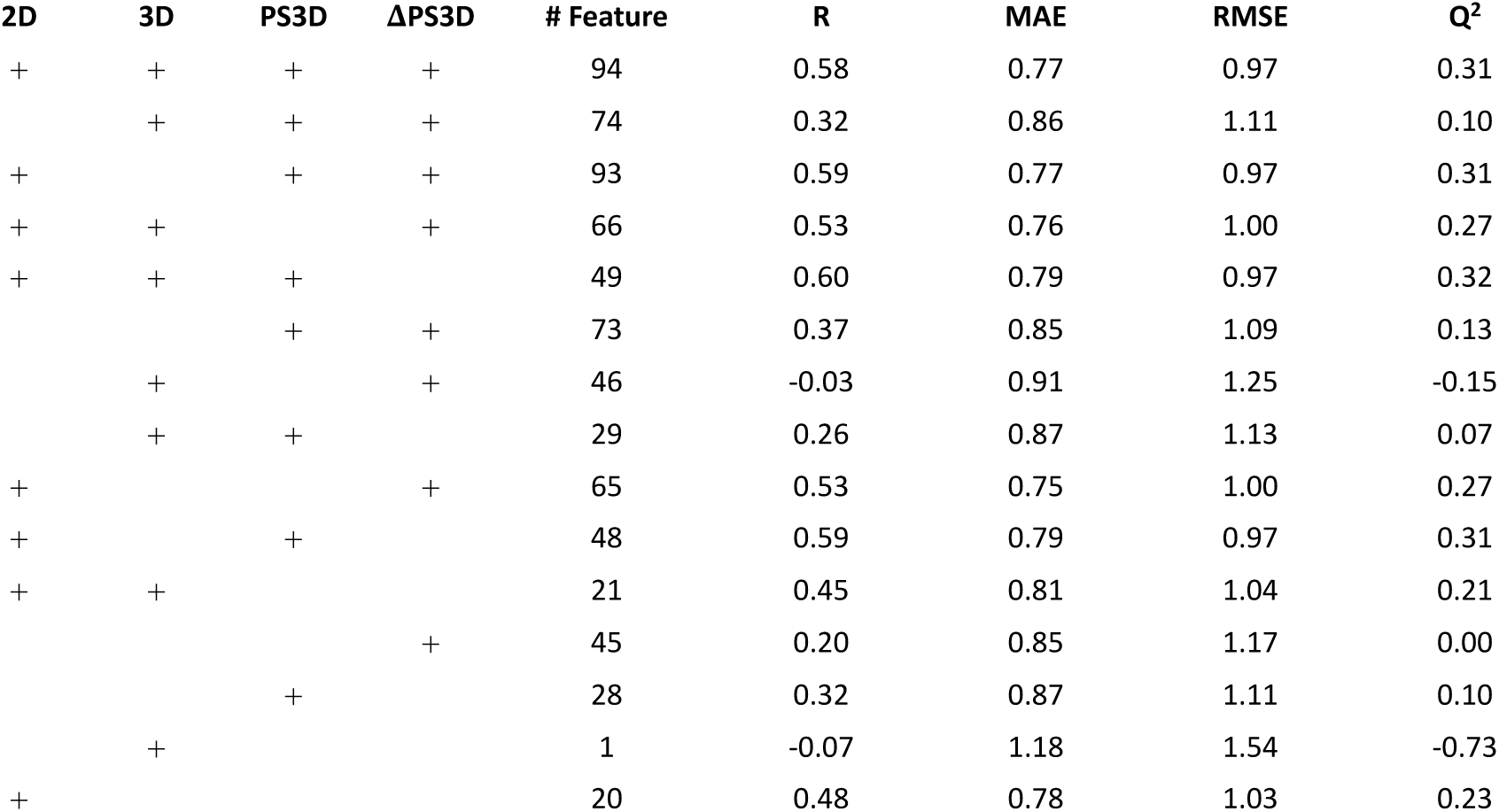
prediction performance when the Bhardwaj2022 was used as test data. A + mark denotes that the corresponding descriptor was used for training and validation.

This study used 252 peptides as the training and validation dataset. However, the number of peptides used was considerably small, and many were synthesized by combinatorically selecting only a few residues from several substituents. Additionally, the dominant chemical species in the dataset did not have a charge near neutral pH, limiting the chemical space covered by our dataset. Consequently, the validation of this study was performed only within a restricted chemical space. However, this study indicated that combining PS3D descriptors and 2D descriptors can predict membrane permeability coefficient, with an average prediction performance of *R =* 0.61, *MAE* = 0.61, *RMSE* = 0.74, and *R*^2^ = 0.28 when one of the four datasets was used as the external test data. Specifically, the results demonstrated that membrane permeability can be predicted with moderate accuracy, even for targets with chemical structure and characteristics of 3D conformation distinct from those of the training data. The proposed method outperforms the use of 2D descriptors alone, particularly in predicting the membrane permeability coefficients of peptides that exist slightly outside of the chemical space covered by the training data. Therefore, the proposed method can predict the membrane permeability of peptides that exist in a wider chemical space by using less training data. In any case, increasing the diversity of training data is expected to increase the model’s universality. This study primarily aims to provide a proof of concept, and larger-scale validation will be performed in the future. Additionally, assessing whether accurately predicted 3D structures improve universality in state-of-the-art deep learning algorithms remains an important challenge for future research. Notably, membrane permeation occurs via various mechanisms.^65^ Particularly, charged peptides—unlike the peptides handled in this study—are thought to permeate membranes other than simple passive diffusion.^66,67^ Therefore, to handle diverse targets, it is crucial to create training datasets tailored to the specific characteristics of those targets.

Since the publication of CycPeptMPDB in 2023, several machine learning models have been proposed to predict membrane permeability of cyclic peptides, including the application of modern deep learning algorithms. ^34–37^ Representative examples are shown in Table 8. The prediction accuracy is variable, indicating that the use of state-of-the-art algorithms does not always result in high prediction accuracy. Notably, PharmPapp, CycPeptMP, and MultiCycPermea:ID reported higher prediction accuracy compared to our validation and external test results (*R*^2^ > 0.65, *MAE* < 0.36). However, the test data used to make those predictions was randomly selected from the entire dataset, with the training data containing peptides from the same literature as the test data. Therefore, their prediction accuracy is overestimated, and there is no guarantee that the predictions for a new dataset will be successful. In contrast, multiCycPermea:OD reported prediction accuracy in a strict setting measured on test data selected, ensuring that Morgan fingerprint was not similar to the training data. The prediction accuracy of MultiCycPermea:OD was slightly lower than the average performance shown in our Table 4–7 which used an arbitrary dataset as test data. This result may suggest an advantage to our method. However, as shown in Tables 2 and 4–7, each data set presents unique challenges in predicting membrane permeability, and the accuracy of prediction ultimately depends on the difficulty of the test data. Therefore, although measuring the prediction accuracy of the entire test data is crucial, it is necessary to focus on the characteristics of each sub-dataset to determine whether the prediction for new data will be successful.

**Table 8.**
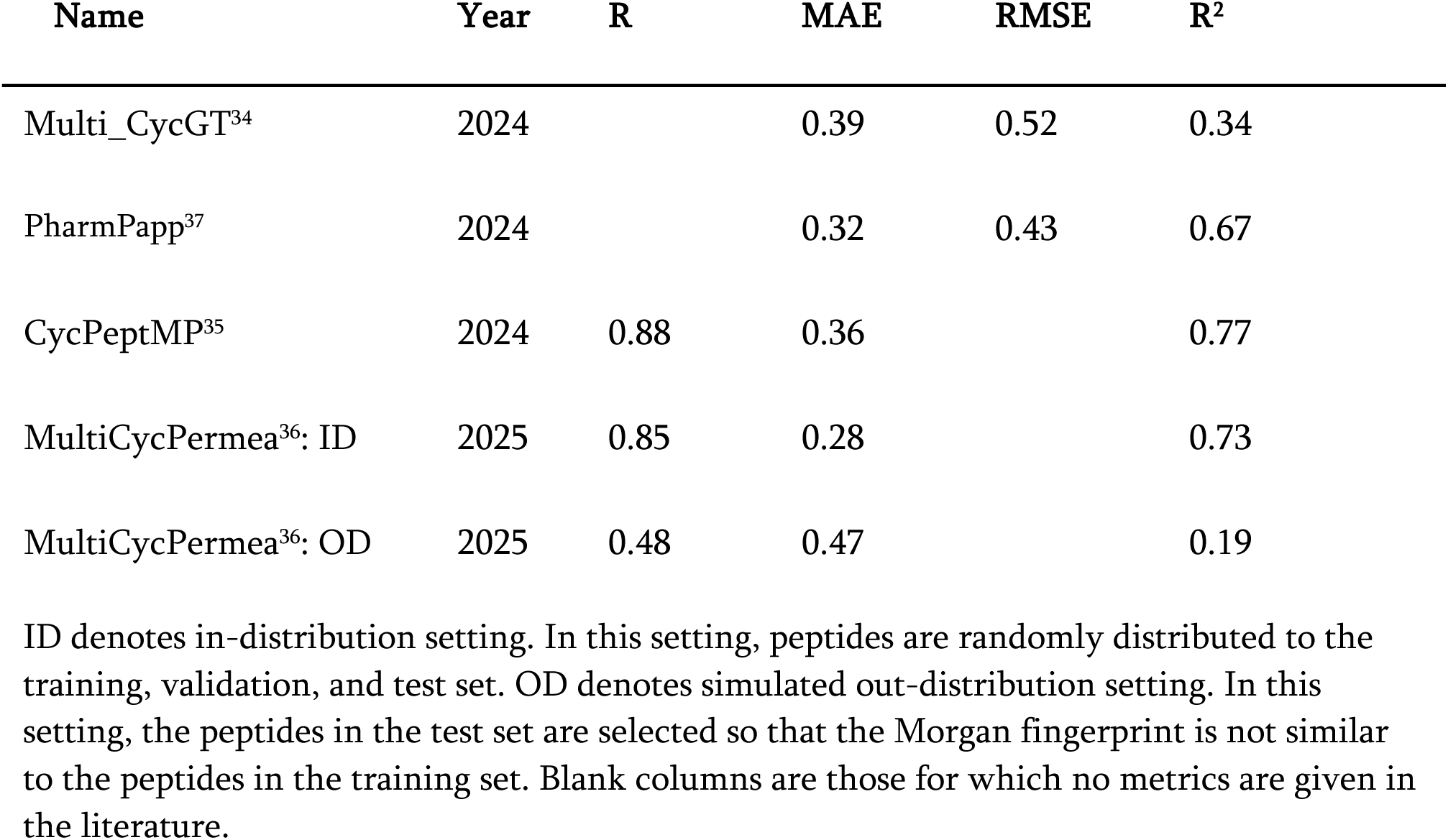
Prediction accuracy of membrane permeability prediction model based on deep learning algorithms using CycPeptMPDB data.

| Name | Year | R | MAE | RMSE | R<sup>2</sup> |
| --- | --- | --- | --- | --- | --- |
| Multi_CycGT<sup>34</sup> | 2024 |  | 0.39 | 0.52 | 0.34 |
| PharmPapp<sup>37</sup> | 2024 |  | 0.32 | 0.43 | 0.67 |
| CycPeptMP<sup>35</sup> | 2024 | 0.88 | 0.36 |  | 0.77 |
| MultiCycPermea<sup>36</sup>: ID | 2025 | 0.85 | 0.28 |  | 0.73 |
| MultiCycPermea<sup>36</sup>: OD | 2025 | 0.48 | 0.47 |  | 0.19 |

ID denotes in-distribution setting. In this setting, peptides are randomly distributed to the training, validation, and test set. OD denotes simulated out-distribution setting. In this setting, the peptides in the test set are selected so that the Morgan fingerprint is not similar to the peptides in the training set. Blank columns are those for which no metrics are given in the literature.

The following outlines the expected procedure for using this method in drug discovery, specifically when applying it to new data. As a preliminary step, it is necessary to accumulate descriptor data (including PS3D descriptors, of course) for training. Although it is recommended to accumulate descriptor data for as many peptides as possible, it is more important to obtain data for peptides that are chemically similar to the prediction target. Subsequently, a subset of this accumulated data should be selected as training data in accordance with the properties of the chemical space of the targets. For example, if one wants to predict the membrane permeability of a group of peptides with similar logP values, it is essential to use such data as training data. Importantly, using the entire peptides in the database as training data is not necessarily optimal.

Even if the number of training data is increased significantly, there is a limit to the accuracy of the prediction. The main reason is that experimental values vary depending on the experimental conditions. For example, in the datasets used in this study, the lower limit of the membrane permeability differs by more than one order of magnitude among the datasets due to differences in experimental conditions, such as assay time and initial concentration, rather than peptide characteristics. Furthermore, when examining the CycPeptMPDB ^33^ for experimental values of peptides such as cyclosporine A, the membrane permeability coefficients obtained by the same assay method for the same peptide vary by more than an order of magnitude across the literature. Therefore, as the number of datasets increases, the diversity of experimental conditions increases, and datasets with similar peptides but different trends in membrane permeability can coexist. This sets an upper limit on prediction accuracy and considerably discourages the use of machine learning algorithms.^68^ To mitigate this issue and develop models with high prediction accuracy, it is necessary to use a large data set, measured under as uniform conditions as possible, as the training data. However, the CycPeptMPDB^34^ contains only a few references with more than 100 experimental values in a single reference. Although new large-scale experimental data has been reported,^16^ it is currently difficult to obtain membrane permeability data for various peptides measured under uniform experimental conditions. As more large-scale experimental data becomes available, it will be possible to develop models with higher prediction accuracy.

Ethical considerations in applying machine learning to drug discovery should be addressed. Prediction results from machine learning algorithms are highly dependent on the quality and characteristics of the training data collected. If unverified data sources are used, the prediction results cannot be considered reliable. In this study, all data were obtained from peer-reviewed papers, ensuring a certain level of reliability. Additionally, if data published by a particular company is used as the primary training data, there is a risk of developing a model that favors peptides only synthesized by that company. This study avoids such a problem by including peptides synthesized by three different research institutions.

## CONCLUSION

In this study, we combined MD simulations and machine learning to predict the membrane permeability coefficient to establish a universally applicable method for predicting membrane permeability at a realistic computational cost. The differences of the prediction quality among the learning algorithms were not large, but when using XGBoost, which showed the best values, we obtained a prediction performance of *R* = 0.77, *MAE* = 0.46, *RMSE* = 0.62, and *Q*^2^ = 0.58. The important descriptors included those that describe the hydrophilicity and hydrophobicity of the peptide, molecular weight, conformational differences between inside and outside the membrane, the degree of freedom of the peptide. When the test data was randomly selected from all datasets, the membrane permeability could be predicted even with only 2D descriptors; however, the values assumed that there were peptides in the training data that were similar to the validation data. To confirm the generalization performance of the method, one of the four datasets was completely removed from the training data and used as test data. The combination of descriptors that produces good prediction results is 2D, and PS3D descriptors, and the average value of the correlation coefficient was *R =* 0.61, *MAE* = 0.61, *RMSE* = 0.74, and *R*^2^ = 0.28. Thus, accurate predictions were not possible using only 2D descriptors and 3D descriptors are essential in practical situations, that is, predicting membrane permeability of peptides that are not similar to the training data. Increasing the diversity of the training data is expected to increase the universality of the model.

Many pharmaceutical companies now routinely perform free energy perturbation (FEP) calculations to predict the binding affinities of small molecule drugs to their targets.^69^ We aimed to reduce the prediction cost of membrane permeability of cyclic peptides to the level of FEP calculations. The ability to accurately predict the membrane permeability of dozens or hundreds of cyclic peptides routinely will facilitate the development of therapeutics for undruggable targets such as intracellular PPI inhibition, a key objective in cyclic peptide drug discovery.

## CONFLICTS OF INTEREST

The authors declare no conflicts of interest associated with this manuscript.

## DATA and SOFTWARE AVAILABILITY

Parameterization, initial conformational sampling, and calculation of descriptors of peptides were performed using MOE (2019.0102, and 2022.02),^53,56^ https://www.chemcomp.com/index.htm. All molecular dynamics simulations and analysis of the trajectories were performed using AMBER 20 and AMBER22 software package,^51,70^ https://ambermd.org/. Python 3.9.6 and the following libraries were used to analyze the results and for machine learning: pandas 2.1.1, numpy 1.24.3, scipy 1.10.1, scikit-learn 1.4.2, xgboost 2.0.3, lightgbm 4.5.0.

## SUPPORTING INFORMATION

The following files are available free of charge.

Detailed protocol for obtaining initial coordinates for the replica exchange with solute tempering/replica exchange umbrella sampling (REST/REUS) simulations; Descriptions of descriptors obtained using CPPTRAJ; 3D descriptor names computed by using MOE; 2D descriptor names computed using MOE; Distribution of the number of residues, molecular weight, and experimental membrane permeability (log scaled) of the dataset; Mean and standard deviation of MAE of predicted and experimental permeability coefficient using random forest algorithm when the threshold of correlation coefficient for reducing the number of descriptors was changed from 0.4 to 0.95 in intervals of 0.05; Number of descriptors used for train and test when the threshold of correlation coefficient for reducing the number of descriptors was changed; Searched hyperparameters for LASSO; Searched hyperparameters for Ridge; Searched hyperparameters for SVM; Searched hyperparameters for RF; Searched hyperparameters for XGBoost; Searched hyperparameters for Light-GBM; Selected optimal hyperparameters for LASSO; Prediction performance of LASSO against each dataset; Prediction performance of Ridge against each dataset; Prediction performance of SVM against each dataset; Prediction performance of RF against each dataset; Prediction performance of Light-GBM against each dataset; Details of 10 descriptors with the highest importance (pdf)

## Author Contributions

M.S. conceived the concept and designed the simulations, performed a part of simulation and analyzed data. Y.N. performed the simulations and analyzed data. J.L. supervised and performed ML part. T.F. analyzed a part of data. K.Y. and Y.A supervised the project. The manuscript was written through contributions of all authors. All authors have given approval to the final version of the manuscript.

## Funding Sources

This work was partially supported by Platform Project for Supporting Drug Discovery and Life Science Research (Basis for Supporting Innovative Drug Discovery and Life Science Research (BINDS)) (Grant No. JP22ama121026) of the Japan Agency for Medical Research and Development (AMED), and FOREST Program (Grant No. JPMJFR216J) of the Japan Science and Technology Agency (JST), KAKENHI (Grant No. 24K15162) of the Japan Society for the Promotion of Science (JSPS).

## ACKNOWLEDGMENT

The numerical calculations were conducted using the TSUBAME 3.0 supercomputer at Tokyo Institute of Technology and TSUBAME 4.0 at Institute of Science Tokyo. A part of the computational resources was awarded by the TSUBAME Grand Challenge Program (spring 2022, spring 2023) from the Global Scientific Information Center, Tokyo Institute of Technology. We would like to thank SAKURA Internet Inc. for collaborative research projects on HPC applications for drug discovery and bioinformatics. We would like to thank Editage (www.editage.jp) for English language editing.

## ABBREVIATIONS

RMSE: root mean square error
MD: molecular dynamics
PAMPA: parallel artificial membrane permeability assay
REST: replica exchange with solute tempering
REUS: replica exchange umbrella sampling method
ST: solute tempering
MOE: molecular operating environment
PS: position-specific
LASSO: LASSO regression
Ridge: ridge regression
SVM: support vector machine
RF: random forest

## Notes

### Competing Interest Statement

The authors have declared no competing interest.

## REFERENCES

(1) Zorzi, A.; Deyle, K.; Heinis, C. Cyclic peptide therapeutics: Past, present and future. Current Opinion in Chemical Biology 2017, 38, 24–29. DOI: 10.1016/j.cbpa.2017.02.006.

(2) Zhang, H.; Chen, S. Cyclic peptide drugs approved in the last two decades (2001−2021). RSC Chemical Biology 2022, 3 (1), 18–31. DOI: 10.1039/d1cb00154j.

(3) Jiang, Y.; Long, H.; Zhu, Y.; Zeng, Y. Macrocyclic peptides as regulators of protein-protein interactions. Chinese Chemical Letters 2018, 29 (7), 1067–1073. DOI: 10.1016/j.cclet.2018.05.028.

(4) Nielsen, D. S.; Shepherd, N. E.; Xu, W.; Lucke, A. J.; Stoermer, M. J.; Fairlie, D. P. Orally absorbed cyclic peptides. Chemical Reviews. 2017, 117 (12), 8094–8128. DOI: 10.1021/acs.chemrev.6b00838.

(5) David J. Craik. Seamless Proteins Tie Up Their Loose Ends. Science (1979) 2006, 311 (5767), 1563–1564.

(6) Goto, Y.; Suga, H. The RaPID platform for the discovery of pseudo-natural macrocyclic peptides. Accounts of Chemical Research 2021, 54 (18), 3604–3617. DOI: 10.1021/acs.accounts.1c00391.

(7) Huang, Y.; Wiedmann, M. M.; Suga, H. RNA display methods for the discovery of bioactive macrocycles. Chemical Reviews 2019, 119 (17) 10360–10391. DOI: 10.1021/acs.chemrev.8b00430.

(8) Bhardwaj, G.; O’Connor, J.; Rettie, S.; Huang, Y. H.; Ramelot, T. A.; Mulligan, V. K.; Alpkilic, G. G.; Palmer, J.; Bera, A. K.; Bick, M. J.; Di Piazza, M.; Li, X.; Hosseinzadeh, P.; Craven, T. W.; Tejero, R.; Lauko, A.; Choi, R.; Glynn, C.; Dong, L.; Griffin, R.; van Voorhis, W. C.; Rodriguez, J.; Stewart, L.; Montelione, G. T.; Craik, D.; Baker, D. Accurate de Novo design of membrane-traversing macrocycles. Cell 2022, 185 (19), 3520–3532.e26. DOI: 10.1016/j.cell.2022.07.019.

(9) Fukunaga, I.; Matsukiyo, Y.; Kaitoh, K.; Yamanishi, Y. Automatic generation of functional peptides with desired bioactivity and membrane permeability using Bayesian Optimization. Molecular Informatics 2024, 43 (4). DOI: 10.1002/minf.202300148.

(10) Ottaviani, G.; Martel, S.; Carrupt, P.-A. Parallel artificial membrane permeability assay: A new membrane for the fast prediction of passive human skin permeability. Journal of Medicinal Chemistry 2006, 49 (13), 3948–3954. DOI: 10.1021/jm060230.

(11) Teixidó, M.; Zurita, E.; Malakoutikhah, M.; Tarragó, T.; Giralt, E. Diketopiperazines as a tool for the study of transport across the Blood-Brain Barrier (BBB) and their potential use as BBB-Shuttles. Journal of the American Chemical Society 2007, 129 (38), 11802–11813. DOI: 10.1021/ja073522o.

(12) Bennion, B. J.; Be, N. A.; McNerney, M. W.; Lao, V.; Carlson, E. M.; Valdez, C. A.; Malfatti, M. A.; Enright, H. A.; Nguyen, T. H.; Lightstone, F. C.; Carpenter, T. S. Predicting a drug’s membrane permeability: A computational model validated with in vitro permeability assay data. Journal of Physical Chemistry B 2017, 121 (20), 5228–5237. DOI: 10.1021/acs.jpcb.7b02914.

(13) Dickson, C. J.; Hornak, V.; Pearlstein, R. A.; Duca, J. S. Structure-kinetic relationships of passive membrane permeation from multiscale modeling. Journal of the American Chemical Society 2017, 139 (1), 442–452. DOI: 10.1021/jacs.6b11215.

(14) Dickson, C. J.; Hornak, V.; Bednarczyk, D.; Duca, J. S. Using membrane partitioning simulations to predict permeability of forty-nine drug-like molecules. Journal of Chemical Information and Modeling 2019, 59 (1), 236–244. DOI: 10.1021/acs.jcim.8b00744.

(15) Danelius, E.; Poongavanam, V.; Peintner, S.; Wieske, L. H. E.; Erdélyi, M.; Kihlberg, J. Solution conformations explain the chameleonic behaviour of macrocyclic drugs. Chemistry - A European Journal 2020, 26 (23), 5231–5244. DOI: 10.1002/chem.201905599.

(16) Faris, J. H.; Adaligil, E.; Popovych, N.; Ono, S.; Takahashi, M.; Nguyen, H.; Plise, E.; Taechalertpaisarn, J.; Lee, H. W.; Koehler, M. F. T.; Cunningham, C. N.; Lokey, R. S. Membrane permeability in a large macrocyclic peptide driven by a saddle-shaped conformation. Journal of the American Chemical Society 2024, 146 (7), 4582–4591. DOI: 10.1021/jacs.3c10949.

(17) Thorpe, M. P.; Hopkins, C. R.; Johnston, J. N. End-to-End backbone cyclization enhances passive permeability of BRo5 Oligomeric Depsipeptides with nonlinear size dependence. ACS Medicinal Chemistry Letters 2025. DOI: 10.1021/acsmedchemlett.5c00037.

(18) Rezai, T.; Bock, J. E.; Zhou, M. V.; Kalyanaraman, C.; Lokey, R. S.; Jacobson, M. P. Conformational flexibility, internal hydrogen bonding, and passive membrane permeability: Successful in silico prediction of the relative permeabilities of cyclic peptides. Journal of the American Chemical Society 2006, 128 (43), 14073–14080. DOI: 10.1021/ja063076p.

(19) Whitty, A.; Zhong, M.; Viarengo, L.; Beglov, D.; Hall, D. R.; Vajda, S. Quantifying the chameleonic properties of macrocycles and other high-molecular-weight drugs. Drug Discovery Today 2016, 21 (5), 712–717. DOI: 10.1016/j.drudis.2016.02.005.

(20) Leung, S. S. F.; Sindhikara, D.; Jacobson, M. P. Simple predictive models of passive membrane permeability incorporating size-dependent membrane-water partition. Journal of Chemical Information and Modeling 2016, 56 (5), 924–929. DOI: 10.1021/acs.jcim.6b00005.

(21) Naylor, M. R.; Ly, A. M.; Handford, M. J.; Ramos, D. P.; Pye, C. R.; Furukawa, A.; Klein, V. G.; Noland, R. P.; Edmondson, Q.; Turmon, A. C.; Hewitt, W. M.; Schwochert, J.; Townsend, C. E.; Kelly, C. N.; Blanco, M. J.; Lokey, R. S. Lipophilic permeability efficiency reconciles the opposing roles of lipophilicity in membrane permeability and aqueous solubility. Journal of Medicinal Chemistry 2018, 61 (24), 11169–11182. DOI: 10.1021/acs.jmedchem.8b01259.

(22) Furukawa, A.; Townsend, C. E.; Schwochert, J.; Pye, C. R.; Bednarek, M. A.; Lokey, R. S. Passive membrane permeability in cyclic peptomer scaffolds is robust to extensive variation in side chain functionality and backbone geometry. Journal of Medicinal Chemistry 2016, 59 (20), 9503–9512. DOI: 10.1021/acs.jmedchem.6b01246.

(23) Ghose, A. K.; Crippen, G. M. Atomic physicochemical parameters for three-dimensional structure-direct Ed quantitative structure-activity relationships I. Partition coefficients as a measure of hydrophobicity. Journal of Computational Chemistry 1986, 7 (4), 565–577.

(24) Sugita, M.; Fujie, T.; Yanagisawa, K.; Ohue, M.; Akiyama, Y. Lipid composition is critical for accurate membrane permeability prediction of cyclic peptides by molecular dynamics simulations. Journal of Chemical Information and Modeling 2022, 62 (18), 4549–4560. DOI: 10.1021/acs.jcim.2c00931.

(25) Loosli, H. -R; Kessler, H.; Oschkinat, H.; Weber, H. -P; Petcher, T. J.; Widmer, A. Peptide conformations. Part 31. The conformation of cyclosporin a in the crystal and in solution. Helvetica Chimica Acta 1985, 68 (3), 682–704. DOI: 10.1002/hlca.19850680319.

(26) Fouché, M.; Schäfer, M.; Berghausen, J.; Desrayaud, S.; Blatter, M.; Piéchon, P.; Dix, I.; Martingarcia, A.; Roth, H. J. Design and development of a cyclic decapeptide scaffold with suitable properties for bioavailability and oral exposure. ChemMedChem 2016, 11 (10), 1048–1059. DOI: 10.1002/cmdc.201600082.

(27) Kelly, C. N.; Townsend, C. E.; Jain, A. N.; Naylor, M. R.; Pye, C. R.; Schwochert, J.; Lokey, R. S. Geometrically diverse lariat peptide scaffolds reveal an untapped chemical space of high membrane permeability. Journal of the American Chemical Society 2021, 143 (2), 705–714. DOI: 10.1021/jacs.0c06115.

(28) Sugita, M.; Sugiyama, S.; Fujie, T.; Yoshikawa, Y.; Yanagisawa, K.; Ohue, M.; Akiyama, Y. Large-scale membrane permeability prediction of cyclic peptides crossing a lipid bilayer based on enhanced sampling molecular dynamics simulations. Journal of Chemical Information and Modeling 2021, 61 (7), 3681–3695. DOI: 10.1021/acs.jcim.1c00380.

(29) Linker, S. M.; Schellhaas, C.; Kamenik, A. S.; Veldhuizen, M. M.; Waibl, F.; Roth, H. J.; Fouché, M.; Rodde, S.; Riniker, S. Lessons for oral bioavailability: How conformationally flexible cyclic peptides enter and cross lipid membranes. Journal of Medicinal Chemistry 2023, 66 (4), 2773–2788. DOI: 10.1021/acs.jmedchem.2c01837.

(30) Linker, S. M.; Schellhaas, C.; Ries, B.; Roth, H. J.; Fouché, M.; Rodde, S.; Riniker, S. Polar/Apolar interfaces modulate the conformational behavior of cyclic peptides with impact on their passive membrane permeability. RSC Advances 2022, 12 (10), 5782–5796. DOI: 10.1039/d1ra09025a.

(31) Witek, J.; Wang, S.; Schroeder, B.; Lingwood, R.; Dounas, A.; Roth, H. J.; Fouché, M.; Blatter, M.; Lemke, O.; Keller, B.; Riniker, S. Rationalization of the membrane permeability differences in a series of analogue cyclic decapeptides. Journal of Chemical Information and Modeling 2019, 59 (1), 294–308. DOI: 10.1021/acs.jcim.8b00485.

(32) Ono, S.; Naylor, M. R.; Townsend, C. E.; Okumura, C.; Okada, O.; Lokey, R. S. Conformation and permeability: Cyclic hexapeptide diastereomers. Journal of Chemical Information and Modeling 2019, 59 (6), 2952–2963. DOI: 10.1021/acs.jcim.9b00217.

(33) Li, J.; Yanagisawa, K.; Sugita, M.; Fujie, T.; Ohue, M.; Akiyama, Y. CycPeptMPDB: A Comprehensive database of membrane permeability of cyclic peptides. Journal of Chemical Information and Modeling 2023, 63 (7), 2240–2250. DOI: 10.1021/acs.jcim.2c01573.

(34) Cao, L.; Xu, Z.; Shang, T.; Zhang, C.; Wu, X.; Wu, Y.; Zhai, S.; Zhan, Z.; Duan, H. Multi_CycGT: A deep learning-based multimodal model for predicting the membrane permeability of cyclic peptides. Journal of Medicinal Chemistry 2024, 67 (3), 1888–1899. DOI: 10.1021/acs.jmedchem.3c01611.

(35) Li, J.; Yanagisawa, K.; Akiyama, Y. CycPeptMP: Enhancing membrane permeability prediction of cyclic peptides with multi-level molecular features and data augmentation. Briefings in Bioinformatics 2024, 25 (5). DOI: 10.1093/bib/bbae417.

(36) Wang, Z.; Chen, Y.; Shang, Y.; Yang, X.; Pan, W.; Ye, X.; Sakurai, T.; Zeng, X. MultiCycPermea: Accurate and interpretable prediction of cyclic peptide permeability using a multimodal image-sequence model. BioMed Central Biology 2025, 23 (1). DOI: 10.1186/s12915-025-02166-2.

(37) Tan, X.; Liu, Q.; Fang, Y.; Zhu, Y.; Chen, F.; Zeng, W.; Ouyang, D.; Dong, J. Predicting peptide permeability across diverse barriers: A systematic investigation. Molecular Pharmaceutics 2024, 21 (8), 4116–4127. DOI: 10.1021/acs.molpharmaceut.4c00478.

(38) Sun, H.; Nguyen, K.; Kerns, E.; Yan, Z.; Yu, K. R.; Shah, P.; Jadhav, A.; Xu, X. Highly predictive and interpretable models for PAMPA permeability. Bioorganic & Medicinal Chemistry 2017, 25 (3), 1266–1276. DOI: 10.1016/j.bmc.2016.12.049.

(39) Rácz, A.; Vincze, A.; Volk, B.; Balogh, G. T. Extending the limitations in the prediction of PAMPA permeability with machine learning algorithms. European Journal of Pharmaceutical Sciences 2023, 188, 106514. DOI: 10.1016/j.ejps.2023.106514.

(40) Lomize, A. L.; Pogozheva, I. D. Physics-based method for modeling passive membrane permeability and translocation pathways of bioactive molecules. Journal of Chemical Information and Modeling 2019, 59 (7), 3198–3213. DOI: 10.1021/acs.jcim.9b00224.

(41) Rossi Sebastiano, M.; Doak, B. C.; Backlund, M.; Poongavanam, V.; Over, B.; Ermondi, G.; Caron, G.; Matsson, P.; Kihlberg, J. Impact of dynamically exposed polarity on permeability and solubility of chameleonic drugs beyond the Rule of 5. Journal of Medicinal Chemistry 2018, 61 (9), 4189–4202. DOI: 10.1021/acs.jmedchem.8b00347.

(42) Kansy, M.; Senner, F.; Gubernator, K. Physicochemical high throughput screening: Parallel artificial membrane permeation assay in the description of passive absorption processes. Journal of Medicinal Chemistry 1998, 41 (7), 1007–1010.

(43) Bermejo, M.; Avdeef, A.; Ruiz, A.; Nalda, R.; Ruell, J. A.; Tsinman, O.; González, I.; Fernández, C.; Sánchez, G.; Garrigues, T. M.; Merino, V. PAMPA - A drug absorption in Vitro Model: 7. Comparing Rat in Situ, Caco-2, and PAMPA permeability of fluoroquinolones. European Journal of Pharmaceutical Sciences 2004, 21 (4), 429–441. DOI: 10.1016/j.ejps.2003.10.009.

(44) Furukawa, A.; Schwochert, J.; Pye, C. R.; Asano, D.; Edmondson, Q. D.; Turmon, A. C.; Klein, V. G.; Ono, S.; Okada, O.; Lokey, R. S. Drug-like properties in macrocycles above MW 1000: Backbone rigidity versus side-chain lipophilicity. Angewandte Chemie - International Edition 2020, 59 (48), 21571–21577. DOI: 10.1002/anie.202004550.

(45) Wang, S.; König, G.; Roth, H. J.; Fouché, M.; Rodde, S.; Riniker, S. Effect of flexibility, lipophilicity, and the location of polar residues on the passive membrane permeability of a series of cyclic decapeptides. Journal of Medicinal Chemistry 2021, 64 (17), 12761– 12773. DOI: 10.1021/acs.jmedchem.1c00775.

(46) Wang, C. K.; Swedberg, J. E.; Harvey, P. J.; Kaas, Q.; Craik, D. J. Conformational flexibility is a determinant of permeability for cyclosporin. Journal of Physical Chemistry B 2018, 122 (8), 2261–2276. DOI: 10.1021/acs.jpcb.7b12419.

(47) Cipcigan, F.; Smith, P.; Crain, J.; Hogner, A.; De Maria, L.; Llinas, A.; Ratkova, E. Membrane permeability in cyclic peptides is modulated by core conformations. Journal of Chemical Information and Modeling 2021, 61 (1), 263–269. DOI: 10.1021/acs.jcim.0c00803.

(48) Jo, S.; Kim, T.; Iyer, V. G.; Im, W. CHARMM-GUI: A web-based graphical user interface for CHARMM. Journal of Computational Chemistry 2008, 29 (11), 1859–1865. DOI: 10.1002/jcc.20945.

(49) Wang, L.; Friesner, R. A.; Berne, B. J. Replica exchange with solute scaling: A more efficient version of replica exchange with solute tempering (REST2). Journal of Physical Chemistry B 2011, 115 (30), 9431–9438. DOI: 10.1021/jp204407d.

(50) Sugita, Y.; Kitao, A.; Okamoto, Y. Multidimensional replica-exchange method for free-energy calculations. Journal of Chemical Physics 2000, 113 (15), 6042–6051. DOI: 10.1063/1.1308516.

(51) Case, D. A.; Belfon, K.; Ben-Shalom, I. Y.; Brozell, S. R.; Cerutti, D. S.; Cheatham, T. E. , III.; Cruzeiro, V. W. D.; Darden, T. A.; Duke, R. E.; Giambasu, G.; Gilson, M. K.; Gohlke, H.; Goetz, A. W.; Harris, R.; Izadi, S.; Izmailov, S. A.; Kasavajhala, K.; Kovalenko, A.; Krasny, R.; Kurtzman, T.; Lee, T. S.; LeGrand, S.; Li, P.; Lin, C.; Liu, J.; Luchkov, T.; Luo, R.; Man, V.; Merz, K. M.; Miao, Y.; Mikhailovskii, O.; Monard, G.; Nguyen, H.; Onufriev, A.; Pan, F.; Pantano, S.; Qi, R.; Roe, D. R.; Roitberg, A.; Sagui, C.; Schott-Verdugo, S.; Shen, J.; Simmerling, C. L.; Skrynnikov, N. R.; Smith, J.; Swails, J.; Walker, R. C.; Wang, J.; Wilson, L.; Wolf, R. M.; Wu, X.; Xiong, Y.; Xue, Y.; York, D. M.; Kollman, P. A. Amber 20. University of California: San Francisco, CA, 2020.

(52) Jorgensen, W. L.; Chandrasekhar, J.; Madura, J. D.; Impey, R. W.; Klein, M. L. Comparison of simple potential functions for simulating liquid water. The Journal of Chemical Physics 1983, 79 (2), 926.

(53) Molecular Operating Environment (MOE), 2019.01; Chemical Computing Group ULC: Canada, 2019.

(54) Ono, S.; Naylor, M. R.; Townsend, C. E.; Okumura, C.; Okada, O.; Lee, H. W.; Lokey, R. S. Cyclosporin A: Conformational complexity and chameleonicity. Journal of Chemical Information and Modeling 2021, 61 (11), 5601–5613. DOI: 10.1021/acs.jcim.1c00771.

(55) Boku, T.; Sugita, M.; Kobayashi, R.; Furuya, S.; Fujie, T.; Ohue, M.; Akiyama, Y. Improving Performance on Replica-Exchange Molecular Dynamics Simulations by Optimizing GPU Core Utilization, Proceedings of the 53rd International Conference on Parallel Processing, Gotland, Sweden, August 12–15, 2024; pp 1082–1091. DOI: 10.1145/3673038.3673097.

(56) Molecular Operating Environment (MOE), 2022.02; Chemical Computing Group ULC: Canada, 2022.

(57) Tibshirani, R. Regression shrinkage and selection via the lasso. Journal of the Royal Statistical Society: Series B (Methodological) 1996, 58 (1), 267–288. DOI: 10.1111/j.2517-6161.1996.tb02080.x.

(58) Hoerl, A. E.; Kennard, R. W. Ridge regression: Biased estimation for nonorthogonal problems. Technometrics 1970, 12 (1), 55–67. DOI: 10.1080/00401706.1970.10488634.

(59) Cortes, C.; Vapnik, V.; Saitta, L. Support-vector networks editor. Machine Leaming 1995, 20, 273–297. DOI: 10.1007/bf00994018.

(60) Breiman, L. Random forests. Machine Learning 2001, 45, 5–32. DOI: 10.1023/a:1010933404324.

(61) Chen, T.; Guestrin, C. XGBoost: A scalable tree boosting system. arXiv:1603.02754 2016. DOI: 10.1145/2939672.2939785.

(62) Ke, G.; Meng, Q.; Finley, T.; Wang, T.; Chen, W.; Ma, W.; Ye, Q.; Liu, T.-Y. LightGBM: A highly efficient gradient boosting decision Tree. Advances in Neural Information Processing Systems 30 (NIPS 2017); 2017.

(63) Boldini, D.; Grisoni, F.; Kuhn, D.; Friedrich, L.; Sieber, S. A. Practical guidelines for the use of gradient boosting for molecular property prediction. Journal of Cheminformatics 2023, 15 (1). DOI: 10.1186/s13321-023-00743-7.

(64) Avdeef, A.; Nielsen, P. E.; Tsinman, O. PAMPA - A drug absorption in vitro model: 11. matching the in vivo unstirred water layer thickness by individual-well stirring in microtitre plates. European Journal of Pharmaceutical Sciences 2004, 22 (5), 365–374. DOI: 10.1016/j.ejps.2004.04.009.

(65) Dougherty, P. G.; Sahni, A.; Pei, D. Understanding cell penetration of cyclic peptides. Chemical Review 2019, 119 (17), 10241–10287. DOI: 10.1021/acs.chemrev.9b00008.

(66) Omatsu, T.; Hori, K.; Ishida, N.; Maeda, K.; Yoshida, Y. Distribution of ion pairs into a bilayer lipid membrane and its effect on the ionic permeability. Biochimica et Biophysica Acta (BBA) - Biomembranes 2021, 1863 (11). DOI: 10.1016/j.bbamem.2021.183724.

(67) Herce, H. D.; Garcia, A. E.; Cardoso, M. C. Fundamental molecular mechanism for the cellular uptake of guanidinium-rich molecules. Journal of the American Chemical Society 2014, 136 (50), 17459–17467. DOI: 10.1021/ja507790z.

(68) Landrum, G. A.; Riniker, S. Combining IC50 or Ki values from different sources is a source of significant noise. Journal of Chemical Information and Modeling 2024, 64 (5), 1560–1567. DOI: 10.1021/acs.jcim.4c00049.

(69) Wang, L.; Wu, Y.; Deng, Y.; Kim, B.; Pierce, L.; Krilov, G.; Lupyan, D.; Robinson, S.; Dahlgren, M. K.; Greenwood, J.; Romero, D. L.; Masse, C.; Knight, J. L.; Steinbrecher, T.; Beuming, T.; Damm, W.; Harder, E.; Sherman, W.; Brewer, M.; Wester, R.; Murcko, M.; Frye, L.; Farid, R.; Lin, T.; Mobley, D. L.; Jorgensen, W. L.; Berne, B. J.; Friesner, R. A.; Abel, R. Accurate and reliable prediction of relative ligand binding potency in prospective drug discovery by way of a modern free-energy calculation protocol and force field. Journal of the American Chemical Society 2015, 137 (*7*), 2695–2703. DOI: 10.1021/ja512751q.

(70) Case, D. A.; Aktulga, H. M.; Belfon, K.; Ben-Shalom, I. Y.; Berryman, J. T.; Brozell, S. ; Cerutti, D. S.; Cheatham, I. T. E.; Cisneros, G. A.; Cruzeiro, V. W. D.; Darden, T. A.; Duke, R. E.; Giambasu, G.; Gilson, M. K.; Gohlke, H.; Goetz, A. W.; Harris, R.; Izadi, ; Izmailov, S. A.; Kasavajhala, K.; Kaymak, M. C.; King, E.; Kovalenko, A.; Kurtzman, S. ; Lee, T. S.; LeGrand, S.; Li, P.; Lin, C.; Liu, J.; Luchko, T.; Luo, R.; Machado, M.; Man, V.; Manathunga, M.; Merz, K. M.; Miao, Y.; Mikhailovskii, O.; Monard, G.; Nguyen, H.; O’Hearn, K. A.; Onufriev, A.; Pan, F.; Pantano, S.; Qi, R.; Rahnamoun, A.; Roe, D. R.; Roitberg, A.; Sagui, C.; Schott-Verdugo, S.; Shajan, A.; Shen, J.; Simmerling, C. L.; Skrynnikov, N. R.; Smith, J.; Swails, J.; Walker, R. C.; Wang, J.; Wang, J.; Wei, H.; Wolf, R. M.; Wu, X.; Xiong, Y.; Xue, Y.; York, D. M.; Zhao, S.; Kollman, P. A. AMBER 22. University of California: San Francisco, CA, 2022. https://ambermd.org/contributors.html.

