## Supplemental material for "Protocol for Membrane Permeability Prediction of Cyclic Peptides using Descriptors Obtained from Extended Ensemble Molecular Dynamics Simulations and Chemical Structures"

**Detailed protocol for obtaining initial coordinates for the replica exchange with solute tempering/replica exchange umbrella sampling (REST/REUS) simulations**

At first, an arbitrary structure of the peptide was placed at $z$ = 40.0 Å, and the structure was minimized for 10,000 steps. In this process, the peptide and lipid molecules were restrained with a harmonic potential, at a 2 kcal/mol/Å$^2$ force constant. The systems were then heated from 0 to 100 K within 20 ps using constant-volume Langevin dynamics, with the restraint of the position of peptide and lipid molecules by harmonic potential, the force constant of which was 1 kcal/mol/Å$^2$. Thereafter, the temperature was increased to 300 K within 100 ps in the isothermal-isobaric (NPT) ensemble with semi-isotropic pressure scaling. The pressure was controlled using a Berendsen barostat and was maintained at 1 bar. The force constant of the harmonic potential for the restrained peptide and lipid molecules decreased to 0.1 kcal/mol/Å$^2$. The systems were equilibrated for 10 ns with positional restraint at $z$ = 40.0 Å. Subsequently, peptides were pulled based on steered MD [1] from $z$ = 40.0 Å to -5.0 Å. A pulling rate of 0.25 Å/ns and force constant of 3.0 kcal/mol/Å$^2$ were used. This steered MD was combined with the ST method and the temperature of the peptide was set to 2100 K. The snapshots for 8 replicas at certain restraint position are extracted from the trajectory of the steered MD located at ± 0.4 Å from the restraint center, which was randomly selected from the trajectory.

**Descriptions of descriptors obtained using CPPTRAJ**

**dE_vdw**

van der Waals interaction energy between a peptide and other molecules.

**dE_elec**

Electrostatic interaction energy between a peptide and other molecules.

**dE**

dE_vdw + dE_elec

**inter_hbond**

Number of hydrogen bonds formed by cyclic peptides and surrounding water molecules. We used a default value for the angle and distance cutoff, i.e., angle > 135° and distance < 3.0 Å.

**intra_hbond**

Number of hydrogen bonds in a cyclic peptide molecule.

**unsatisfied_hb_donor**

Number of hydrogen bonded donors without intramolecular hydrogen bonding, i.e., number of hydrogen bonded donors minus intra_hbond value.

**polar_surface_area**

Accessible surface area of nitrogen and oxygen atoms.

**entropy**

Estimated conformation entropy based on quasi-harmonic analysis.[2]

**dihedral_pca_cos**

Cosine similarity between the conformation distributions projected onto the first and second principal components obtained from principal component analysis of the three main chain angles (φ, ψ, Ω).

**dihedral_pca_emd**

Earth mover's distance [3] between the distributions of conformations projected onto the first and second principal components obtained from principal component analysis of the three main chain angles (φ, ψ, Ω).

**p_a_i_r**

A numerical value representing the rough shape of a molecule calculated based on equation S1. [4] Note that the moments of inertia denote Ia, Ib, and Ic in ascending order from the smallest eigenvalue.

$$r = \frac{2\sqrt{\frac{1}{I_b} + \frac{1}{I_c}}}{\sqrt{\frac{1}{I_a} + \frac{1}{I_b}} + \sqrt{\frac{1}{I_a} + \frac{1}{I_c}}} \quad (S1)$$

**p_a_i_angle_1**

Angle between the principal axis of inertia corresponding to the smallest eigenvalue and reaction coordinate z.

**p_a_i_angle_2**

Angle between the principal axis of inertia corresponding to the intermediate eigenvalue and reaction coordinate z.

**p_a_i_angle_3**

Angle between the principal axis of inertia corresponding to the largest eigenvalue and reaction coordinate z.

**LW_contacts**

Number of interactions between the carbon atoms of cholesterol, palmitic acid, and oleic acid and oxygen, sodium, and chlorine atoms of the solvent molecule. The cutoff value to determine interaction was set to 4 Å.

**membrane_size**

Average volume of the space comprising the membrane. The average area of the x-y plane of the simulation box multiplied by the average distance of the centers of mass of the nitrogen atoms in the POPC head between the upper and lower leaflets.

**Table S1.**

| Dataset | Usage for Traning/validation or test | Number of peptides selected for the data set | Number of peptides reported in the reference | Referenced paper |
|---|---|---|---|---|
| Furukawa2016 | traning/validation | 67 | 688 | Furukawa et al., *J. Med. Chem.*, 2016 [5] |
| Furukawa2020 | traning/validation | 18 | 36 | Furukawa et al., *Angew. Chem. Int. Ed.*, 2020 [6] |
| Kelly2021 | traning/validation | 86 | 1519 | Kelly et al., *J. Am. Chem. Soc.*, 2021 [7] |
| Bhardwaj2022 | traning/validation | 81 | 136 | Bhardwaj et al., *Cell*, 2022 [8] |
| Wang2021 | test | 24 | 24 | Wang et al., *J. Med. Chem.*, 2021 [9] |

**Table S2**. 3D descriptor names computed using MOE

| index | name | description |
|---|---|---|
| 1 | ASA | Water accessible surface area calculated using a radius of 1.4 Å for the water molecule. |
| 2 | ASA+ | Water accessible surface area of all atoms with positive partial charge. |
| 3 | ASA− | Water accessible surface area of all atoms with negative partial charge. |
| 4 | ASA_H | Water accessible surface area of all hydrophobic atoms. |
| 5 | ASA_P | Water accessible surface area of all polar atoms. |
| 6 | CASA+ | Positive charge weighted surface area, ASA+ times max { qi > 0 }. |
| 7 | CASA− | Negative charge weighted surface area, ASA- times max { qi < 0 } . |
| 8 | DASA | Absolute value of the difference between ASA+ and ASA-. |
| 9 | DCASA | Absolute value of the difference between CASA+ and CASA-. |
| 10 | E | Value of the potential energy. |
| 11 | E_ang | Angle bend potential energy. |
| 12 | E_ele | Electrostatic component of the potential energy. |
| 13 | E_nb | Value of the potential energy with all bonded terms disabled. |
| 14 | E_oop | Out-of-plane potential energy. |
| 15 | E_sol | Solvation energy. |
| 16 | E_stb | Bond stretch-bend cross-term potential energy. |
| 17 | E_str | Bond stretch potential energy. |
| 18 | E_strain | The current energy minus the value of the energy at a near local minimum. |
| 19 | E_tor | Torsion potential energy. |
| 20 | E_vdw | van der Waals component of the potential energy. |
| 21 | FASA+ | Fractional ASA+ calculated as ASA+ / ASA. |
| 22 | FASA− | Fractional ASA- calculated as ASA- / ASA. |
| 23 | FASA_H | Fractional ASA_H calculated as ASA_H / ASA. |
| 24 | FASA_P | Fractional ASA_P calculated as ASA_P / ASA. |
| 25 | FCASA+ | Fractional CASA+ calculated as CASA+ / ASA. |
| 26 | FCASA− | Fractional CASA- calculated as CASA- / ASA. |
| 27 | VSA | van der Waals surface area. |
| 28 | dens | Molecular weight divided by van der Waals volume as calculated in the vol descriptor. |
| 29 | dipole | Dipole moment calculated from the partial charges of the molecule. |
| 30 | glob | Globularity, or inverse condition number (smallest eigenvalue divided by the largest eigenvalue) of the covariance matrix of atomic coordinates. |

**Table S3**. 3D descriptor names computed using MOE (Cont'd).

| index | name | description |
|---|---|---|
| 31 | npr1 | Normalized PMI ratio pmi1/pmi3. |
| 32 | npr2 | Normalized PMI ratio pmi2/pmi3. |
| 33 | pmi | Principal moment of inertia. |
| 34 | pmi1 | First diagonal element of diagonalized moment of inertia tensor. |
| 35 | pmi2 | Second diagonal element of diagonalized moment of inertia tensor. |
| 36 | pmi3 | Third diagonal element of diagonalized moment of inertia tensor. |
| 37 | rgyr | Radius of gyration. |
| 38 | std_dim1 | The square root of the largest eigenvalue of the covariance matrix of the atomic coordinates. |
| 39 | std_dim2 | The square root of the second largest eigenvalue of the covariance matrix of the atomic coordinates. |
| 40 | std_dim3 | The square root of the third largest eigenvalue of the covariance matrix of the atomic coordinates. |
| 41 | vol | van der Waals volume calculated using a grid approximation (spacing 0.75 A). |
| 42 | vsurf_A | Amphiphilic moment |
| 43 | vsurf_CP | Critical packing parameter |
| 44 | vsurf_CW1 | Capacity factor at -0.2 |
| 45 | vsurf_CW2 | Capacity factor at -0.5 |
| 46 | vsurf_CW3 | Capacity factor at -1.0 |
| 47 | vsurf_CW4 | Capacity factor at -2.0 |
| 48 | vsurf_CW5 | Capacity factor at -3.0 |
| 49 | vsurf_CW6 | Capacity factor at -4.0 |
| 50 | vsurf_CW7 | Capacity factor at -5.0 |
| 51 | vsurf_CW8 | Capacity factor at -6.0 |
| 52 | vsurf_D1 | Hydrophobic volume at -0.2 |
| 53 | vsurf_D2 | Hydrophobic volume at -0.4 |
| 54 | vsurf_D3 | Hydrophobic volume at -0.6 |
| 55 | vsurf_D4 | Hydrophobic volume at -0.8 |
| 56 | vsurf_D5 | Hydrophobic volume at -1.0 |
| 57 | vsurf_D6 | Hydrophobic volume at -1.2 |
| 58 | vsurf_D7 | Hydrophobic volume at -1.4 |
| 59 | vsurf_D8 | Hydrophobic volume at -1.6 |
| 60 | vsurf_DD12 | vsurf_EDmin1, vsurf_EDmin2 distance |

**Table S4**. 3D descriptor names computed using MOE (Cont'd).

| index | name | index |
|---|---|---|
| 61 | vsurf_DD13 | vsurf_EDmin1, vsurf_EDmin3 distance |
| 62 | vsurf_DD23 | vsurf_EDmin2, vsurf_EDmin3 distance |
| 63 | vsurf_DW12 | vsurf_EWmin1, vsurf_EWmin2 distance |
| 64 | vsurf_DW13 | vsurf_EWmin1, vsurf_EWmin3 distance |
| 65 | vsurf_DW23 | vsurf_EWmin2, vsurf_EWmin3 distance |
| 66 | vsurf_EDmin1 | Lowest hydrophobic energy |
| 67 | vsurf_EDmin2 | 2nd lowest hydrophobic energy |
| 68 | vsurf_EDmin3 | 3rd lowest hydrophobic energy |
| 69 | vsurf_EWmin1 | Lowest hydrophilic energy |
| 70 | vsurf_EWmin2 | 2nd lowest hydrophilic energy |
| 71 | vsurf_EWmin3 | 3rd lowest hydrophilic energy |
| 72 | vsurf_G | Surface globularity |
| 73 | vsurf_HB1 | H-bond donor capacity at -0.2 |
| 74 | vsurf_HB2 | H-bond donor capacity at -0.5 |
| 75 | vsurf_HB3 | H-bond donor capacity at -1.0 |
| 76 | vsurf_HB4 | H-bond donor capacity at -2.0 |
| 77 | vsurf_HB5 | H-bond donor capacity at -3.0 |
| 78 | vsurf_HB6 | H-bond donor capacity at -4.0 |
| 79 | vsurf_HB7 | H-bond donor capacity at -5.0 |
| 80 | vsurf_HB8 | H-bond donor capacity at -6.0 |
| 81 | vsurf_HL1 | First hydrophilic-lipophilic balance |
| 82 | vsurf_HL2 | Second hydrophilic-lipophilic balance |
| 83 | vsurf_ID1 | Hydrophobic integy moment at -0.2 |
| 84 | vsurf_ID2 | Hydrophobic integy moment at -0.4 |
| 85 | vsurf_ID3 | Hydrophobic integy moment at -0.6 |
| 86 | vsurf_ID4 | Hydrophobic integy moment at -0.8 |
| 87 | vsurf_ID5 | Hydrophobic integy moment at -1.0 |
| 88 | vsurf_ID6 | Hydrophobic integy moment at -1.2 |
| 89 | vsurf_ID7 | Hydrophobic integy moment at -1.4 |
| 90 | vsurf_ID8 | Hydrophobic integy moment at -1.6 |

**Table S5**. 3D descriptor names computed using MOE (Cont'd).

| index | name | description |
|---|---|---|
| 91 | vsurf_IW1 | Hydrophilic integy moment at -0.2 |
| 92 | vsurf_IW2 | Hydrophilic integy moment at -0.5 |
| 93 | vsurf_IW3 | Hydrophilic integy moment at -1.0 |
| 94 | vsurf_IW4 | Hydrophilic integy moment at -2.0 |
| 95 | vsurf_IW5 | Hydrophilic integy moment at -3.0 |
| 96 | vsurf_IW6 | Hydrophilic integy moment at -4.0 |
| 97 | vsurf_IW7 | Hydrophilic integy moment at -5.0 |
| 98 | vsurf_IW8 | Hydrophilic integy moment at -6.0 |
| 99 | vsurf_R | Surface rugosity |
| 100 | vsurf_S | Interaction field surface area |
| 101 | vsurf_V | Interaction field volume |
| 102 | vsurf_W1 | Hydrophilic volume at -0.2 |
| 103 | vsurf_W2 | Hydrophilic volume at -0.5 |
| 104 | vsurf_W3 | Hydrophilic volume at -1.0 |
| 105 | vsurf_W4 | Hydrophilic volume at -2.0 |
| 106 | vsurf_W5 | Hydrophilic volume at -3.0 |
| 107 | vsurf_W6 | Hydrophilic volume at -4.0 |
| 108 | vsurf_W7 | Hydrophilic volume at -5.0 |
| 109 | vsurf_W8 | Hydrophilic volume at -6.0 |
| 110 | vsurf_Wp1 | Polar volume at -0.2 |
| 111 | vsurf_Wp2 | Polar volume at -0.5 |
| 112 | vsurf_Wp3 | Polar volume at -1.0 |
| 113 | vsurf_Wp4 | Polar volume at -2.0 |
| 114 | vsurf_Wp5 | Polar volume at -3.0 |
| 115 | vsurf_Wp6 | Polar volume at -4.0 |
| 116 | vsurf_Wp7 | Polar volume at -5.0 |
| 117 | vsurf_Wp8 | Polar volume at -6.0 |

**Table S6**. 2D descriptor names computed using MOE

| index | name | description |
|---|---|---|
| 1 | BCUT_PEOE_0 | The smallest eigenvalue of the partial charges (PCs) based on partial equalization of orbital electronegativities (PEOE) method, derived using the the BCUT method. |
| 2 | BCUT_PEOE_1 | The 1/3 percentile eigenvalue of the PEOE PCs based on BCUT method. |
| 3 | BCUT_PEOE_2 | The 2/3 percentile eigenvalue of the PEOE PCs based on BCUT method. |
| 4 | BCUT_PEOE_3 | The largest percentile eigenvalue of the PEOE PCs based on BCUT method. |
| 5 | BCUT_SLOGP_0 | The smallest percentile eigenvalue of the SlogP based on BCUT method. |
| 6 | BCUT_SLOGP_1 | The 1/3 percentile eigenvalue of the SlogP based on BCUT method. |
| 7 | BCUT_SLOGP_2 | The 2/3 percentile eigenvalue of the SlogP based on BCUT method. |
| 8 | BCUT_SLOGP_3 | The largest percentile eigenvalue of the SlogP based on BCUT method. |
| 9 | BCUT_SMR_0 | The smallest percentile eigenvalue of the SMR based on BCUT method. |
| 10 | BCUT_SMR_1 | The 1/3 percentile eigenvalue of the SMR based on BCUT method. |
| 11 | BCUT_SMR_2 | The 2/3 percentile eigenvalue of the SMR based on BCUT method. |
| 12 | BCUT_SMR_3 | The largest percentile eigenvalue of the SMR based on BCUT method. |
| 13 | GCUT_PEOE_0 | The smallest percentile eigenvalue of the PEOE PCs based on GCUT method. GCUT descriptors are the values obtained by replacing the bond order in BCUT descriptors with distance. |
| 14 | GCUT_PEOE_1 | The 1/3 percentile eigenvalue of the PEOE PCs based on GCUT method. |
| 15 | GCUT_PEOE_2 | The 2/3 percentile eigenvalue of the PEOE PCs based on GCUT method. |
| 16 | GCUT_PEOE_3 | The largest percentile eigenvalue of the PEOE PCs based on GCUT method. |
| 17 | GCUT_SLOGP_0 | The smallest percentile eigenvalue of the SlogP based on GCUT method. |
| 18 | GCUT_SLOGP_1 | The 1/3 percentile eigenvalue of the SlogP based on GCUT method. |
| 19 | GCUT_SLOGP_2 | The 2/3 percentile eigenvalue of the SlogP based on GCUT method. |
| 20 | GCUT_SLOGP_3 | The largest percentile eigenvalue of the SlogP based on GCUT method. |
| 21 | GCUT_SMR_0 | The GCUT descriptors using atomic contribution to molar refractivity (using the Wildman and Crippen SMR method) instead of PC. |
| 22 | GCUT_SMR_1 | The smallest percentile eigenvalue of the SMR based on GCUT method. |
| 23 | GCUT_SMR_2 | The 1/3 percentile eigenvalue of the SMR based on GCUT method. |
| 24 | GCUT_SMR_3 | The 2/3 percentile eigenvalue of the SMR based on GCUT method. |
| 25 | Kier1 | The largest percentile eigenvalue of the SMR based on GCUT method. |
| 26 | Kier2 | Second kappa shape index. |
| 27 | Kier3 | Third kappa shape index. |
| 28 | KierA1 | First alpha modified shape index. |
| 29 | KierA2 | Second alpha modified shape index. |
| 30 | KierA3 | Third alpha modified shape index. |

**Table S7**. 2D descriptor names computed using MOE (Cont'd)

| index | name | description |
|---|---|---|
| 31 | KierFlex | Kier molecular flexibility index. |
| 32 | PC+ | Total positive PC. |
| 33 | PC− | Total negative PC. |
| 34 | PEOE_PC+ | Total positive PEOE PC. |
| 35 | PEOE_PC− | Total negative PEOE PC. |
| 36 | PEOE_RPC+ | Relative positive PEOE PC. |
| 37 | PEOE_RPC− | Relative negative PEOE PC. |
| 38 | PEOE_VSA+0 | Sum of van der Waals (vdW) surface areas (SAs) where PEOE PC is in the range [0.00,0.05). |
| 39 | PEOE_VSA+1 | Sum of vdW SAs where PEOE PC is in the range [0.05,0.10). |
| 40 | PEOE_VSA+2 | Sum of vdW SAs where PEOE PC is in the range [0.10,0.15). |
| 41 | PEOE_VSA+3 | Sum of vdW SAs where PEOE PC is in the range [0.15,0.20). |
| 42 | PEOE_VSA+4 | Sum of vdW SAs where PEOE PC is in the range [0.20,0.25). |
| 43 | PEOE_VSA+5 | Sum of vdW SAs where PEOE PC is in the range [0.25,0.30). |
| 44 | PEOE_VSA−0 | Sum of vdW SAs where PEOE PC is in the range [-0.05,0.00). |
| 45 | PEOE_VSA−1 | Sum of vdW SAs where PEOE PC is in the range [-0.10,-0.05). |
| 46 | PEOE_VSA−3 | Sum of vdW SAs where PEOE PC is in the range [-0.20,-0.15). |
| 47 | PEOE_VSA−4 | Sum of vdW SAs where PEOE PC is in the range [-0.25,-0.20). |
| 48 | PEOE_VSA−5 | Sum of vdW SAs where PEOE PC is in the range [-0.30,-0.25). |
| 49 | PEOE_VSA−6 | Sum of vdW SAs where PEOE PC is less than -0.30. |
| 50 | PEOE_VSA_FHYD | The sum of the vdW SAs such that absolute value of the PEOE PC is less than or equal to 0.2 divided by the total SA. |
| 51 | PEOE_VSA_FNEG | The sum of the vdW SAs such that PEOE PC is negative divided by the total SA. |
| 52 | PEOE_VSA_FPNEG | The sum of the vdW SAs such that PEOE PC is less than -0.2 divided by the total SA. |
| 53 | PEOE_VSA_FPOL | The sum of the vdW SAs such that absolute value of the PEOE PC is greater than 0.2 divided by the total SA. |
| 54 | PEOE_VSA_FPOS | The sum of the vdW SAs such that PEOE PC is non-negative divided by the total SA. |
| 55 | PEOE_VSA_FPPOS | The sum of the vdW SAs such that PEOE PC is greater than 0.2 divided by the total SA. |
| 56 | PEOE_VSA_HYD | The sum of the vdW SAs such that absolute value of the PEOE PC is less than or equal to 0.2. |
| 57 | PEOE_VSA_NEG | The sum of the vdW SAs such that PEOE PC is negative. |
| 58 | PEOE_VSA_PNEG | The sum of the vdW SAs such that PEOE PC is less than -0.2. |

**Table S8**. 2D descriptor names computed using MOE (Cont'd)

| index | name | description |
|---|---|---|
| 59 | PEOE_VSA_POL | The sum of the vdW SAs such that absolute value of the PEOE PC is greater than 0.2. |
| 60 | PEOE_VSA_POS | The sum of the vdW SAs such that PEOE PC is non-negative. |
| 61 | PEOE_VSA_PPOS | The sum of the vdW SAs such that PEOE PC is greater than 0.2. |
| 62 | Q_PC+ | The sum of the positive PCs. |
| 63 | Q_PC− | The sum of the negative PCs. |
| 64 | Q_RPC+ | The largest positive PC divided by the sum of the positive PCs. |
| 65 | Q_RPC− | The smallest negative PC divided by the sum of the negative PCs. |
| 66 | Q_VSA_FHYD | The sum of the vdW SAs such that absolute value of the PC is less than or equal to 0.2 divided by the total SA. |
| 67 | Q_VSA_FNEG | The sum of the vdW SAs such that PC is negative divided by the total SA. |
| 68 | Q_VSA_FPNEG | The sum of the vdW SAs such that PC is less than -0.2 divided by the total SA. |
| 69 | Q_VSA_FPOL | The sum of the vdW SAs such that absolute value of the PC is greater than 0.2 divided by the total SA. |
| 70 | Q_VSA_FPOS | The sum of the vdW SAs such that PC is non-negative divided by the total SA. |
| 71 | Q_VSA_FPPOS | The sum of the vdW SAs such that PC is greater than 0.2 divided by the total SA. |
| 72 | Q_VSA_HYD | The sum of the vdW SAs such that absolute value of the PC is less than or equal to 0.2. |
| 73 | Q_VSA_NEG | Total negative vdW SA. This is the sum of the vdW SA such that PC is negative. |
| 74 | Q_VSA_PNEG | The sum of the vdW SAs such that PC is less than -0.2. |
| 75 | Q_VSA_POL | The sum of the vdW SAs such that absolute value of the PC is greater than 0.2. |
| 76 | Q_VSA_POS | The sum of the vdW SAs such that PC is non-negative. |
| 77 | Q_VSA_PPOS | The sum of the vdW SAs such that PC is greater than 0.2. |
| 78 | RBC | The number of bonds which match a rule-based definition of rotatable bonds. |
| 79 | RPC+ | The largest positive PC divided by the sum of the positive PCs. |
| 80 | RPC− | The smallest negative PC divided by the sum of the negative PCs. |
| 81 | SMR | Molecular refractivity is an atomic contribution model. |
| 82 | SMR_VSA0 | Sum of vdW SAs for atoms where the contribution to SMR is in [0,0.11]. |
| 83 | SMR_VSA1 | Sum of vdW SAs for atoms where the contribution to SMR is in (0.11,0.26]. |
| 84 | SMR_VSA2 | Sum of vdW SAs for atoms where the contribution to SMR is in (0.26,0.35]. |
| 85 | SMR_VSA3 | Sum of vdW SAs for atoms where the contribution to SMR is in (0.35,0.39]. |
| 86 | SMR_VSA4 | Sum of vdW SAs for atoms where the contribution to SMR is in (0.39,0.44]. |
| 87 | SMR_VSA5 | Sum of vdW SAs for atoms where the contribution to SMR is in (0.44,0.485]. |
| 88 | SMR_VSA6 | Sum of vdW SAs for atoms where the contribution to SMR is in (0.485,0.56]. |
| 89 | SMR_VSA7 | Sum of vdW SAs for atoms where the contribution to SMR > 0.56. |

**Table S9**. 2D descriptor names computed using MOE (Cont'd)

| index | name | description |
|---|---|---|
| 90 | SlogP | The log of the octanol/water partition coefficient is an atomic contribution model. |
| 91 | SlogP_VSA0 | Sum of vdW SAs for atoms where the contribution to SlogP <= -0.4. |
| 92 | SlogP_VSA1 | Sum of vdW SAs for atoms where the contribution to SlogP is in (-0.4,-0.2]. |
| 93 | SlogP_VSA2 | Sum of vdW SAs for atoms where the contribution to SlogP is in (-0.2,0]. |
| 94 | SlogP_VSA3 | Sum of vdW SAs for atoms where the contribution to SlogP is in the range (0,0.1]. |
| 95 | SlogP_VSA4 | Sum of vdW SAs for atoms where the contribution to SlogP is in (0.1,0.15]. |
| 96 | SlogP_VSA5 | Sum of vdW SAs for atoms where the contribution to SlogP is in (0.15,0.20]. |
| 97 | SlogP_VSA6 | Sum of vdW SAs for atoms where the contribution to SlogP is in (0.20,0.25]. |
| 98 | SlogP_VSA7 | Sum of vdW SAs for atoms where the contribution to SlogP is in (0.25,0.30]. |
| 99 | SlogP_VSA8 | Sum of vdW SAs for atoms where the contribution to SlogP is in (0.30,0.40]. |
| 100 | SlogP_VSA9 | Sum of vdW SAs for atoms where the contribution to SlogP > 0.40. |
| 101 | TPSA | Polar SA calculated using group contributions to approximate the polar SA from connection table information only. |
| 102 | VAdjEq | Vertex adjacency information (equality). |
| 103 | VAdjMa | Vertex adjacency information (magnitude). |
| 104 | VDistEq | If m is the sum of the distance matrix entries then VdistEq is defined to be the sum of log2 m - pi log2 pi / m where pi is the number of distance matrix entries equal to i. |
| 105 | VDistMa | If m is the sum of the distance matrix entries then VDistMa is defined to be the sum of log2 m - Dij log2 Dij / m over all i and j. |
| 106 | Weight | Molecular weight (including implicit hydrogens) in atomic mass units with atomic weights. |
| 107 | a_IC | Atom information content (total). This is calculated to be a_ICM times n. |
| 108 | a_ICM | The entropy of the element distribution in the molecule (including implicit hydrogens but not lone pair pseudo-atoms). |
| 109 | a_acc | Number of hydrogen bond acceptor atoms (not counting acidic atoms but counting atoms that are both hydrogen bond donors and acceptors such as -OH). |
| 110 | a_aro | Number of aromatic atoms. |
| 111 | a_count | Number of atoms, including implicit hydrogens. |
| 112 | a_don | Number of hydrogen bond donor atoms (not counting basic atoms but counting atoms that are both hydrogen bond donors and acceptors such as -OH). |
| 113 | a_donacc | Number of hydrogen bond donor plus number of hydrogen bond acceptor atoms. |
| 114 | a_heavy | Number of heavy atoms. |
| 115 | a_hyd | Number of hydrophobic atoms. |
| 116 | a_nC | Number of carbon atoms. |
| 117 | a_nCl | Number of chlorine atoms. |

**Table S10**. 2D descriptor names computed using MOE (Cont'd)

| index | name | description |
|---|---|---|
| 118 | a_nH | Number of hydrogen atoms (including implicit hydrogens). |
| 119 | a_nN | Number of nitrogen atoms. |
| 120 | a_nNO | Number of nitrogen and oxygen atoms. |
| 121 | a_nO | Number of oxygen atoms. |
| 122 | a_nS | Number of sulfur atoms. |
| 123 | apol | Sum of the atomic polarizabilities (including implicit hydrogens) with polarizabilities. |
| 124 | arorings | The number of aromatic rings. |
| 125 | ast_violation | The number of violations of the Astex fragment-like test. |
| 126 | ast_violation_ext | The number of violations of the extended Astex fragment-like test, ast_violation + (opr_nrot > 3) + (vsa_pol > 60). |
| 127 | b_1rotN | The number of rotatable single bonds, excluding conjugated single bonds (e.g. ester and peptide bonds). |
| 128 | b_1rotR | b_1rotN divided by b_heavy. |
| 129 | b_ar | Number of aromatic bonds. |
| 130 | b_count | Number of bonds (including implicit hydrogens). |
| 131 | b_double | Number of double bonds. Aromatic bonds are not considered to be double bonds. |
| 132 | b_heavy | Number of bonds between heavy atoms. |
| 133 | b_max1len | Length of the longest single bond chain. |
| 134 | b_rotN | Number of rotatable bonds, where a bond is rotatable if it has order 1, is not in a ring, and has at least two heavy neighbors. |
| 135 | b_rotR | b_rotN divided by b_heavy. |
| 136 | b_single | Number of single bonds (including implicit hydrogens). Aromatic bonds are not considered to be single bonds. |
| 137 | balabanJ | Balaban's connectivity topological index. |
| 138 | bpol | Sum of the absolute value of the difference between atomic polarizabilities of all bonded atoms in the molecule with polarizabilities. |
| 139 | chi0 | Atomic connectivity index (order 0). |
| 140 | chi0_C | Carbon connectivity index (order 0). |
| 141 | chi0v | Atomic valence connectivity index (order 0). |
| 142 | chi0v_C | Carbon valence connectivity index (order 0). |
| 143 | chi1 | Atomic connectivity index (order 1). |
| 144 | chi1_C | Carbon connectivity index (order 1). This is calculated as the sum of 1/sqrt(didj) over all bonds between carbon atoms i and j where i < j. |
| 145 | chi1v | Atomic valence connectivity index (order 1). |
| 146 | chi1v_C | Carbon valence connectivity index (order 1). |

**Table S11**. 2D descriptor names computed using MOE (Cont'd)

| index | name | description |
|---|---|---|
| 147 | chiral | The number of chiral centers. |
| 148 | density | Weight divided by vdw_vol. |
| 149 | diameter | Largest value in the distance matrix. |
| 150 | h_ema | Sum of hydrogen bond acceptor strengths. |
| 151 | h_emd | Sum of hydrogen bond donor strengths. |
| 152 | h_emd_C | Sum of hydrogen bond donor strengths of carbon atoms. |
| 153 | h_logD | The octanol/water distribution coefficient at pH 7. |
| 154 | h_logP | The logarithm of the octanol/water partition coefficient using an 8 parameter model. |
| 155 | h_logS | The logarithm of the aqueous solubility using a 7 parameter model. |
| 156 | h_log_pbo | The sum of logarithm of one plus the pi bond order for all bonds. |
| 157 | h_mr | The molar refractivity using a 4 parameter model. |
| 158 | h_pKb | The pKb of the reaction that adds a proton from the ensemble of states with a hydrogen count equal to the input structure;14 is reported if there are no states with more hydrogens than the input. |
| 159 | h_pavgQ | The average total charge sum { $Q_i$ $10^{-pC_i}$ } where $Q_i$ is the total formal charge of state i. |
| 160 | h_pstates | The entropic count or fractional number of protonation states. |
| 161 | h_pstrain | The strain energy needed to convert all protonation states into the input protonation state. |
| 162 | lip_acc | The number of O and N atoms. |
| 163 | lip_don | The number of OH and NH atoms. |
| 164 | lip_violation | The number of violations of Lipinski's Rule of Five. |
| 165 | logP(o/w) | The logarithm of the octanol/water partition coefficient is calculated using a linear atom type model. |
| 166 | logS | The logarithm of the aqueous solubility is calculated from an atom contribution linear atom type model. |
| 167 | mr | The molecular refractivity is calculated from an 11 descriptor linear model. |
| 168 | mutagenic | Indicator of the presence of potentially toxic groups. A non-zero value indicates that the molecule contains a mutagenic group.. |
| 169 | opr_brigid | The number of rigid bonds. |
| 170 | opr_nring | The number of rings. |
| 171 | opr_nrot | The number of rotatable bonds. |
| 172 | opr_violation | The number of violations of Oprea's lead-like test. |
| 173 | petitjean | Value of (diameter - radius) / diameter. |

**Table S12**. 2D descriptor names computed using MOE (Cont'd)

| index | name | description |
|---|---|---|
| 174 | petitjeanSC | Petitjean graph Shape Coefficient. |
| 175 | radius | If $r_i$ is the largest matrix entry in row i of the distance matrix D, then the radius is defined as the smallest of the $r_i$. |
| 176 | reactive | Indicator of the presence of reactive groups. A non-zero value indicates that the molecule contains a reactive group. |
| 177 | rings | The number of rings. |
| 178 | rsynth | A value in [0,1] indicating the likelihood of synthesizing the chemical structure, where 0 is unlikely and 1 is likely. |
| 179 | vdw_area | Value of vdW SA. |
| 180 | vdw_vol | Value of vdW volume. |
| 181 | vsa_acc | Approximation to the sum of vdW SAs of pure hydrogen bond acceptors (not counting atoms that are both hydrogen bond donors and acceptors such as -OH). |
| 182 | vsa_don | Approximation to the sum of vdW SAs of pure hydrogen bond donors (not counting atoms that are both hydrogen bond donors and acceptors such as -OH). |
| 183 | vsa_hyd | Approximation to the sum of vdW SAs of hydrophobic atoms. |
| 184 | vsa_other | Approximation to the sum of vdW SAs of atoms typed as "other". |
| 185 | vsa_pol | Approximation to the sum of vdW SAs of polar atoms (atoms that are both hydrogen bond donors and acceptors), such as -OH. |
| 186 | wienerPath | Wiener path number: half the sum of all the distance matrix entries. |
| 187 | wienerPol | Wiener polarity number: half the sum of all the distance matrix entries with a value of 3. |
| 188 | zagreb | The sum of the square of the number of heavy neighbors for all heavy atoms. |

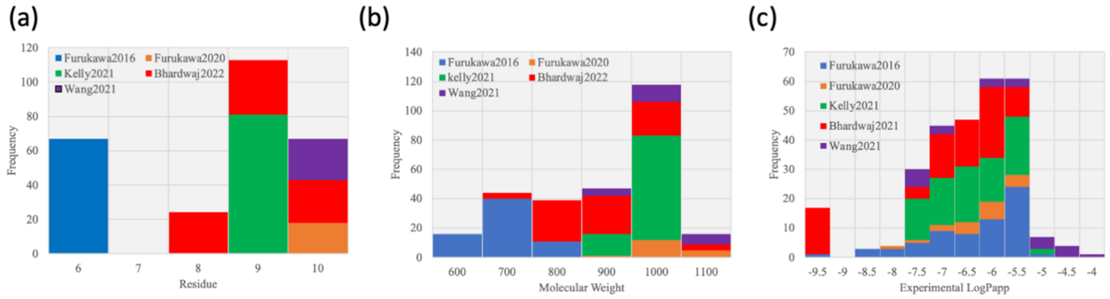


**Figure S1**. Distribution of the number of residues, molecular weight, and experimental membrane permeability (log scaled) of the dataset.

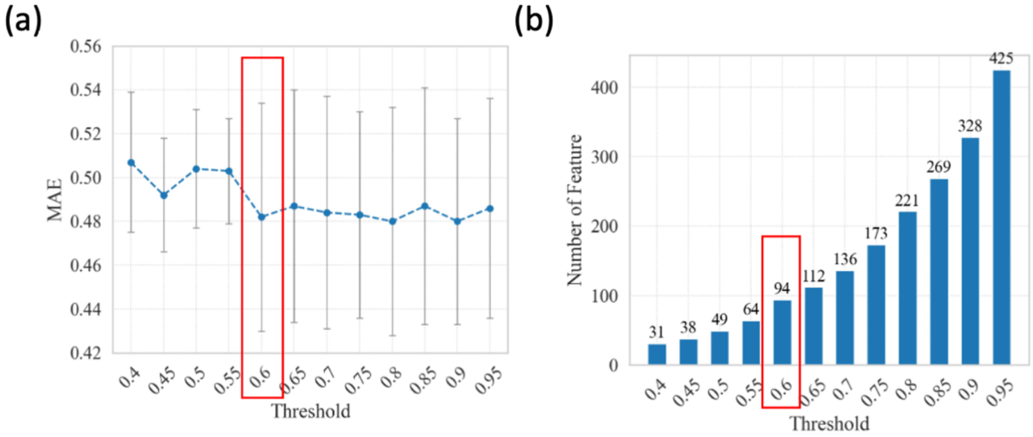


**Figure S2**. (a)Mean and standard deviation of MAE of predicted and experimental permeability coefficient using random forest algorithm when the threshold of correlation coefficient for reducing the number of descriptors was changed from 0.4 to 0.95 in intervals of 0.05. (b) Number of descriptors used for train and test when the threshold of correlation coefficient for reducing the number of descriptors was changed.

**Table S13**. Searched hyperparameter for LASSO.

| parameter | Searched range | Number of cases | Used Value |
|---|---|---|---|
| alpha | $10^{-3}, 10^{-2.9}, \ldots, 10^{1.5}$ | 46 | $10^{-1.5}$ |

**Table S14**. Searched hyperparameter for Ridge.

| parameter | Searched range | Number of cases | Used value |
|---|---|---|---|
| alpha | $10^{-2}, 10^{-1.9}, \ldots, 10^{2.5}$ | 46 | $10^{2.0}$ |

**Table S15**. Searched hyperparameters for support vector machine (SVM).

| parameter | Searched range | Number of cases | Used value |
|---|---|---|---|
| C | $2^{-5}, 2^{-4}, \ldots, 2^{10}$ | 16 | $2^{2}$ |
| epsilon | $2^{-6}, 2^{-5}, \ldots, 2^{0}$ | 7 | $2^{-4}$ |
| gamma | $2^{-15}, 2^{-14}, \ldots, 2^{0}$ | 21 | $2^{-9}$ |

**Table S16**. Searched hyperparameters for random forest (RF).

| parameter | Searched range | Number of cases | Used value |
|---|---|---|---|
| max_depth | None, 2, 5, 10, 15, 20 | 6 | 15 |
| max_features | sqrt, log2, None | 3 | None |
| min_sample_split | $2^{1}, 2^{2}, 2^{3}, 2^{4}$ | 4 | 2 |
| n_estimators | 100, 200, 300, 400, 500, 600, 700 | 7 | 700 |

**Table S17**. Searched hyperparameters for XGBoost.

| parameter | Searched range | Number of cases | Used value |
|---|---|---|---|
| gamma | 0, 0.1, 0.2, 0.3 | 4 | 0 |
| learning_rate | 0.01, 0.1, 0.3 | 3 | 0.3 |
| max_depth | 2, 4, 6, 8 | 4 | 8 |
| min_child_weight | 1, 3, 5 | 3 | 1 |
| n_estimators | 100, 200, 300, 400, 500, 600, 700 | 7 | 700 |
| reg_alpha | 0, 0.1, 0.5, 1 | 4 | 0 |
| reg_lambda | 0, 0.1, 0.5, 1 | 4 | 0 |
| subsample | 0.4, 0.6, 0.8, 1.0 | 4 | 0.4 |

**Table S18**. Searched hyperparameters for Light-GBM.

| parameter | Searched range | Number of cases | Used value |
|---|---|---|---|
| learning_rate | 0.01, 0.1, 0.2, 0.3 | 40 | 0.2 |
| max_depth | -1, 2, 4, 6, 8, 10, 15 | 6 | -1 |
| min_data_in leaf | 5, 10, 15, 20, 30 | 5 | 5 |
| num_leaves | 5, 10, 20, 30, 40 | 5 | 5 |
| n_estimators | 100, 200, 300, 400, 500 | 5 | 100 |
| reg_alpha | 0, 0.1, 0.5, 1.0 | 4 | 0.1 |
| reg_lambda | 0, 0.1, 0.5, 1.0 | 4 | 0.1 |

**Table S19**. Prediction performance of LASSO against each dataset.

| Data | R | RMSE |
|---|---|---|
| Furukawa2016 | $0.81 \pm 0.06$ | $0.52 \pm 0.08$ |
| Furukawa2020 | $0.63 \pm 0.47$ | $0.56 \pm 0.19$ |
| Kelly2021 | $0.85 \pm 0.05$ | $0.43 \pm 0.08$ |
| Bhardwaj2022 | $0.66 \pm 0.13$ | $0.86 \pm 0.12$ |

**Table S20**. Prediction performance of Ridge against each dataset.

| Data | R | RMSE |
|---|---|---|
| Furukawa2016 | 0.77 $\pm$ 0.10 | 0.57 $\pm$ 0.10 |
| Furukawa2020 | 0.66 $\pm$ 0.49 | 0.55 $\pm$ 0.17 |
| Kelly2021 | 0.81 $\pm$ 0.06 | 0.44 $\pm$ 0.07 |
| Bhardwaj2022 | 0.68 $\pm$ 0.11 | 0.83 $\pm$ 0.13 |

**Table S21**. Prediction performance of SVM against each dataset.

| Data | R | RMSE |
|---|---|---|
| Furukawa2016 | 0.80 $\pm$ 0.09 | 0.53 $\pm$ 0.09 |
| Furukawa2020 | 0.70 $\pm$ 0.49 | 0.48 $\pm$ 0.17 |
| Kelly2021 | 0.80 $\pm$ 0.07 | 0.47 $\pm$ 0.07 |
| Bhardwaj2022 | 0.67 $\pm$ 0.11 | 0.83 $\pm$ 0.13 |

**Table S22**. Prediction performance of RF against each dataset.

| Data | R | RMSE |
|---|---|---|
| Furukawa2016 | 0.82 $\pm$ 0.05 | 0.55 $\pm$ 0.07 |
| Furukawa2020 | 0.66 $\pm$ 0.41 | 0.63 $\pm$ 0.28 |
| Kelly2021 | 0.88 $\pm$ 0.05 | 0.38 $\pm$ 0.07 |
| Bhardwaj2022 | 0.64 $\pm$ 0.15 | 0.86 $\pm$ 0.15 |

**Table S23**. Prediction performance of Light-GBM against each dataset.

| Data | R | RMSE |
|---|---|---|
| Furukawa2016 | 0.80 $\pm$ 0.05 | 0.54 $\pm$ 0.08 |
| Furukawa2020 | 0.59 $\pm$ 0.53 | 0.61 $\pm$ 0.32 |
| Kelly2021 | 0.85 $\pm$ 0.04 | 0.42 $\pm$ 0.07 |
| Bhardwaj2022 | 0.59 $\pm$ 0.12 | 0.91 $\pm$ 0.13 |

**Table S24**. Details of 10 descriptors with the highest importance. 2D, 3D, PS3D, and ΔPS3D denotes that the descriptor is 2D, 3D, position specific (PS) 3D, and difference of PS3D descriptor.

| Descriptor | Description |
|---|---|
| SlogP (2D) | Log scaled octanol/water partition coefficient. [10] |
| density (2D) | Molecular mass density: Weight divided by vdW volume (amu/$Å^3$). |
| GCUT_SLOGP_1 (2D) | An eigenvalue of a modified graph distance adjacency matrix.[11] Each ij entry of the adjacency matrix takes the value 1/sqr(dij) where dij is the modified graph distance between atoms i and j. The diagonal takes the value of the atomic contribution to SlogP. |
| vsa_pol (2D) | Approximation to the sum of vdW surface areas ($Å^2$) of polar atoms (atoms that are both hydrogen bond donors and acceptors), such as -OH. |
| opr_nrot (2D) | The number of rotatable bonds.[12] |
| GCUT_PEOE_0 (2D) | An eigenvalue of a modified graph distance adjacency matrix. [11] Each ij entry of the adjacency matrix takes the value 1/sqr(dij) where dij is the modified graph distance between atoms i and j. The diagonal takes the value of the partial charges calculated based on Partial Equalization of Orbital Electronegativities (PEOE) method.[13] |
| E_str (3D) | Bond stretch potential energy. |
| dihedral_pca_cos (membrane − water) (ΔPS3D) | Cosine distance of the distributions of conformations of the peptide between membrane center and in solution. The distribution is projected onto the first and second principal components obtained from principal component analysis of the three main chain angles (φ, ψ, Ω). |
| vsurf_CW5_sur (PS3D) | Capacity factor of the peptide at the water/membrane interface.[14] |
| vsurf_IW1_mem − vsurf_IW1_sur (ΔPS3D) | Difference of hydrophilic integy moment [14] between water/membrane interface and membrane center. |

**References**


(1) Park, S.; Khalili-Araghi, F.; Tajkhorshid, E.; Schulten, K. Free Energy Calculation from Steered Molecular Dynamics Simulations Using Jarzynski's Equality. *J. Chem. Phys.* **2003**, *119* (6), 3559–3566. DOI: 10.1063/1.1590311.

(2) Chang, C. E.; Chen, W.; Gilson, M. K. Evaluating the Accuracy of the Quasiharmonic Approximation. *J. Chem. Theory. Comput.* **2005**, *1* (5), 1017–1028. DOI: 10.1021/ct0500904.

(3) Rubner, Y.; Tomasi, C.; Guibas, L. J. A Metric for Distributions with Applications to Image Databases *. In *Sixth International Conference on Computer Vision (IEEE Cat. No.98CH36271)*. **1998**, 59–66.

(4) Mizuno-Kaneko, M.; Hashimoto, I.; Miyahara, K.; Kochi, M.; Ohashi, N.; Tsumura, K.; Suzuki, K.; Tamura, T. Molecular Design of Cyclic Peptides with Cell Membrane Permeability and Development of MDMX-P53 Inhibitor. *ACS Med. Chem. Lett.* **2023**, *14* (9), 1174–1178. DOI: 10.1021/acsmedchemlett.3c00102.

(5) Furukawa, A.; Townsend, C. E.; Schwochert, J.; Pye, C. R.; Bednarek, M. A.; Lokey, R. S. Passive Membrane Permeability in Cyclic Peptomer Scaffolds Is Robust to Extensive Variation in Side Chain Functionality and Backbone Geometry. *J. Med. Chem.* **2016**, *59* (20), 9503–9512. DOI: 10.1021/acs.jmedchem.6b01246.

(6) Furukawa, A.; Schwochert, J.; Pye, C. R.; Asano, D.; Edmondson, Q. D.; Turmon, A. C.; Klein, V. G.; Ono, S.; Okada, O.; Lokey, R. S. Drug-Like Properties in Macrocycles above MW 1000: Backbone Rigidity versus Side-Chain Lipophilicity. *Angew. Chem., Int. Ed.* **2020**, *59* (48), 21571–21577. DOI: 10.1002/anie.202004550.

(7) Kelly, C. N.; Townsend, C. E.; Jain, A. N.; Naylor, M. R.; Pye, C. R.; Schwochert, J.; Lokey, R. S. Geometrically Diverse Lariat Peptide Scaffolds Reveal an Untapped Chemical Space of High Membrane Permeability. *J. Am. Chem. Soc.* **2021**, *143* (2), 705–714. DOI: 10.1021/jacs.0c06115.

(8) Bhardwaj, G.; O'Connor, J.; Rettie, S.; Huang, Y. H.; Ramelot, T. A.; Mulligan, V. K.; Alpkilic, G. G.; Palmer, J.; Bera, A. K.; Bick, M. J.; Di Piazza,

M.; Li, X.; Hosseinzadeh, P.; Craven, T. W.; Tejero, R.; Lauko, A.; Choi, R.; Glynn, C.; Dong, L.; Griffin, R.; van Voorhis, W. C.; Rodriguez, J.; Stewart, L.; Montelione, G. T.; Craik, D.; Baker, D. Accurate de Novo Design of Membrane-Traversing Macrocycles. *Cell* **2022**, *185* (19), 3520-3532.e26. DOI: 10.1016/j.cell.2022.07.019.

(9) Wang, S.; König, G.; Roth, H. J.; Fouché, M.; Rodde, S.; Riniker, S. Effect of Flexibility, Lipophilicity, and the Location of Polar Residues on the Passive Membrane Permeability of a Series of Cyclic Decapeptides. *J. Med. Chem.* **2021**, *64* (17), 12761–12773. DOI: 10.1021/acs.jmedchem.1c00775.

(10) Wildman, S. A.; Crippen, G. M. Prediction of Physicochemical Parameters by Atomic Contributions. *J. Chem. Inf. Comput. Sci.* **1999**, *39* (5), 868–873. DOI: 10.1021/ci990307l.

(11) Petitjean, M. Applications of the Radius-Diameter Diagram to the Classification of Topological and Geometrical Shapes of Chemical Compounds. *J. Chem. Inf. Comput. Sci.* **1992**, *32* (4), 331–337. DOI: 10.1021/ci00008a012

(12) Oprea, T. I. Property Distribution of Drug-Related Chemical Databases. *J. Comput. Aided. Mol. Des.* **2000**, *14* (3), 251–264. DOI: 10.1023/a:1008130001697.

(13) Gasteiger, J.; Marsili, M. Iterative Partial Equalization of Orbital Electronegativity—a Rapid Access to Atomic Charges. *Tetrahedron* **1980**, *36* (22), 3219–3228. DOI: 10.1016/0040-4020(80)80168-2

(14) Cruciani, G.; Pastor, M.; Guba, W. VolSurf: A New Tool for the Pharmacokinetic Optimization of Lead Compounds. *Eur. J. Pharm. Sci.*, *11* (2), S29–S39. DOI: 10.1016/S0928-0987(00)00162-7
